# Insights from the reanalysis of high-throughput chemical genomics data for *Escherichia coli* K-12

**DOI:** 10.1101/2020.07.16.206243

**Authors:** Peter I-Fan Wu, Curtis Ross, Deborah A. Siegele, James C. Hu

## Abstract

Despite the demonstrated success of genome-wide genetic screens and chemical genomics studies at predicting functions for genes of unknown function or predicting new functions for well-characterized genes, their potential to provide insights into gene function hasn’t been fully explored. We systematically reanalyzed a published high-throughput phenotypic dataset for the model Gram-negative bacterium *Escherichia coli* K-12. The availability of high-quality annotation sets allowed us to compare the power of different metrics for measuring phenotypic profile similarity to correctly infer gene function. We conclude that there is no single best method; the three metrics tested gave comparable results for most gene pairs. We also assessed how converting qualitative phenotypes to discrete, qualitative phenotypes affected the association between phenotype and function. Our results indicate that this approach may allow phenotypic data from different studies to be combined to produce a larger dataset that may reveal functional connections between genes not detected in individual studies.

## INTRODUCTION

Genome-wide genetic screens and chemical genomic studies, pioneered in yeast (GIAEVER AND NISLOW 2014), are now widely used to study gene function in many model organisms, including the bacterium *Escherichia coli* (Campos et al., 2018; Nichols et al., 2011; Price et al., 2018). Based on the same principle that underlies the interpretation of forward genetic studies — that mutations that cause similar phenotypes are likely to affect the same biological process(es) — these high-throughput approaches have led to insights into the biology of a variety of organisms (Arnoldo et al., 2014; Hillenmeyer et al., 2010; Shefchek et al., 2020). It has been concluded that the collective phenotypic expression pattern of an organism can serve as a key to understand growth, fitness, development, and diseases (Bochner, 2009; Houle et al, 2010).

Despite the demonstrated success of high-throughput phenotypic studies at predicting functions for genes of unknown function or predicting new functions for well-characterized genes, their potential to provide insights into gene function hasn’t been fully explored. There does not seem to have been a systematic comparison of different metrics for measuring the similarity of phenotypic profiles. Further, while the likely benefits of combining information from high throughput phenotypic studies from different laboratories have been recognized, very few methods of doing this have been described (Hoehndorf et al., 2013; Shefchek et al., 2020).

Here, we report reanalysis of the data from a published high-throughput phenotypic study of *Escherichia coli* K-12 (Nichols et al. 2011). *E. coli* is one of the best-studied bacterial organisms, and the availability of high-quality annotation sets with information on gene function and regulation allowed us to compare the ability of different metrics for measuring phenotypic profile similarity to correctly infer gene function. We conclude that there is no single best method for comparing phenotypic profiles. Overall, the three metrics we tested gave comparable results for most gene pairs. However, there were instances where the metrics behaved differently from one another. We also assessed how converting quantitative phenotypes to discrete, qualitative phenotypes affected associations between phenotype and function. Our results indicate that this may be a viable approach for combining phenotypic data from different studies, creating a larger dataset that may reveal functional associations not detected by individual studies alone.

## RESULTS

### Phenotypic profiles and the functional annotation sets used

We start with descriptions of the phenotype data and functional annotation sets that were used for our analysis. The phenotypic profiles come from a high-throughput chemical genomics study of *E. coli* K-12 (Nichols et al., 2011). Growth phenotypes for 3,979 mutant strains, which were primarily single-gene deletions of non-essential genes, were based on sizes of spot colonies grown under 324 conditions, which represented 114 unique stresses. Fitness scores were obtained and normalized to a standard normal distribution based on the mean fitness for all strains in a given condition. Positive scores indicate increased fitness and negative scores indicate decreases fitness. Fitness scores were obtained and normalized to a standard normal distribution where positive scores indicate increased fitness and negative scores indicate decreased fitness, which was based on the mean fitness for all strains in a given growth condition.

Six annotation sets were used as sources of information about gene function. Annotations of *E. coli* genes to metabolic pathways and protein complexes were obtained from EcoCyc (Keseler et al., 2017); annotation of genes to operons and regulons were extracted from EcoCyc and RegulonDB (Gama-Castro et al., 2016); and annotations of genes to KEGG modules, which associate genes to metabolic pathways, molecular complexes, and also to phenotypic groups, such as pathogenesis or drug resistance, were obtained from the Kyoto Encyclopedia of Genes and Genomes (KEGG) (Kanehisa et al, 2016). For these annotation sets, genes were scored as co-annotated if they shared the same annotation(s) from one or more of the annotation sets, for example, being annotated to the same metabolic pathway or protein complex, etc. The number of genes annotated by each annotation set and the total number of annotations can be found in Materials and Methods.

The annotations of *E. coli* genes with Gene Ontology (GO) biological process terms (Gene Ontology Consortium, 2017) were obtained from EcoCyc. The GO biological process annotations of *E. coli* genes were treated separately from the other five annotation sets because GO’s directed-acyclic graph structure allows semantic similarity rather than co-annotation to be used for assessing functional similarity (Pesquita, 2017). While it is possible to identify gene pairs that are co-annotated with the same GO term(s), automated methods will include co-annotations to high-level terms, such as ‘GO:0044237 cellular metabolic process’ or ‘GO:0051716 cellular response to stimulus’, which don’t provide very specific information about function. Also, co-annotation doesn’t capture instances where two genes are annotated with related, but not identical, terms. These limitation can be overcome by using semantic similarity rather than co-annotation to estimate functional similarity from GO annotations. The method for determining the semantic similarity of two GO terms developed by Wang et al. (Wang et al, 2007), takes into account the locations of the terms in the GO graph, as well as incorporating the different semantic contributions that a shared ancestral term may make to the two terms, based on the logical relationship, such as is_a or part_of, that connect the term to the shared ancestor. In addition, when calculating functional similarity, the Wang method includes both identical GO terms and semantically similar GO terms associated with the two genes being compared. The number of genes annotated with GO biological process terms set and the total number of annotations can be found in the Materials and Methods.

### Functional connections between genes enriched for higher phenotypic profile similarity

The association between phenotypic profiles and functional annotations was examined from two perspectives: First, are gene pairs that share the same annotation(s), i.e. co-annotated gene pairs, more likely to have higher phenotypic profile similarity? Second, are gene pairs with higher phenotypic profile similarity more likely to be co-annotated?

To address whether co-annotated gene pairs have higher phenotypic profile similarity, we used Pearson Correlation Coefficient (PCC) to assess the phenotypic profile similarity. This metric was chosen because it is probably the most widely used metric to assess phenotypic profile similarity and was the metric used in the original paper for comparing phenotypic profiles (Nichols et al., 2011). To visualize the results, the distributions of the absolute value of PCC (|PCC|) for gene pairs were plotted as violin plots for various combinations of annotation sets (Figure 1). The first violin plot shows the distribution of |PCC| values for all possible gene pairs (mean |PCC| = 0.00016). The majority have a |PCC| value <0.25 and only 0.16% have a |PCC| value >0.75 (an arbitrarily chosen cut-off). When only gene pairs that are co-annotated to the same EcoCyc pathway were considered (second violin plot), there was a statistically significant increase in the mean |PCC| value (0.032), and the percentage of gene pairs with |PCC| >0.75 increased twenty-fold. Similar results were seen for gene pairs that are co-annotated to the same heteromeric protein complex (third violin plot, mean |PCC| = 0.05). When considering only gene pairs that are co-annotated to more than one annotation set (fourth and fifth violin plots), even higher phenotypic profile similarity was observed (mean |PCC| = 0.19, 0.30, respectively), supporting the expectation that gene pairs with stronger functional associations will have more similar phenotypic profiles. The trend of there being a higher fraction of gene pairs with |PCC| >0.75 as functional associations increase also continued; this fraction increased from 0.16% for all gene pairs, to 3.2% for gene pairs in the same pathways, to 4.9% for gene pairs in the same protein complexes, to 19% for gene pairs in the same pathways and complexes, and to 30% for gene pairs that are co-annotated in pathways, complexes, operons, regulons and KEGG modules.

**Figure 1.**
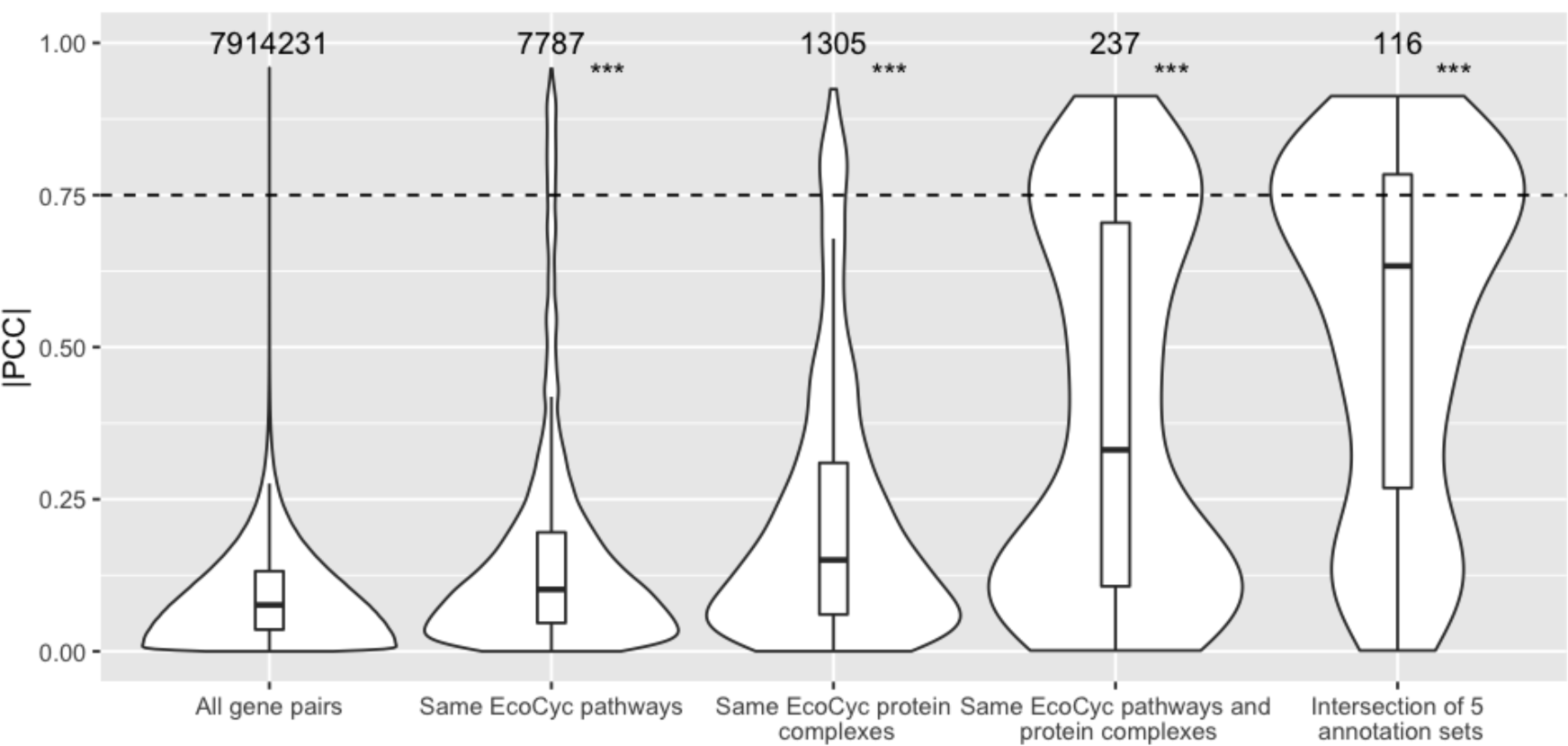
Higher phenotypic similarity was found for co-annotated gene pairs. Shown are violin plots of the distributions of |PCC| for the indicated groups of gene pairs. Numbers above each violin plot indicate the number of gene pairs in each plot. ***: p-value <0.001 was determined by 1-sided Mann-Whitney U test, compared to all gene pairs. The dashed line indicates |PCC| = 0.75, which was chosen as an arbitrary cut-off.

A more detailed analysis within the EcoCyc pathway or heteromeric protein complex annotations was conducted by examining all pairwise combinations of gene pairs within pathways or protein complexes that contain two or more gene products. Supplemental Figures S1 and S2 show the distribution of |PCC| values for all pairwise combinations of genes in each pathway or protein complex. Of the 366 pathways and 271 protein complexes analyzed, 72% of the pathways and 67% of the protein complexes had a median |PCC| value that was higher than the random expectation.

### Phenotypic profile similarity is explained by functional annotations

To address the second question, which is to test whether gene pairs with higher phenotypic profile similarity are more likely to be co-annotated, we ranked gene pairs based on phenotypic profile similarity and then calculated precision based on whether or not gene pairs are co-annotated (Figure 2). Precision is the fraction of results that a test identifies as positive that represent true positives. Mathematically, precision, also known as the positive predictive value, is the number of True Positives divided by True Positives plus False Positives, or TP/(TP+FP). After ranking gene pairs based on phenotypic profile similarity expressed as |PCC| values, precision for each position *n* in the ranking was calculated considering gene pairs ranked at or above position *n* to be TPs if they are co-annotated or FPs if they are not co-annotated. For example, for the 100th gene pair in the ranking, precision is calculated for gene pairs 1 through 100. Figure 2 shows the plots of precision versus ranking for the top-ranking 500 gene pairs computed for single annotation sets or combinations of annotation sets. For gene pairs co-annotated to the same pathway(s), precision started at zero, because the highest ranked gene pair was not co-annotated, but then increased to ∼0.8 before gradually declining and leveling off at approximately 0.2. Surprisingly, for gene pairs co-annotated to the same protein complex, precision was very low and not significantly different from the precision values computed for randomly ordered gene pairs. Combining the annotation sets for pathways and protein complexes, brought a slight increase in precision. When operon, regulon, and KEGG modules were also included to define the broadest set of co-annotations, precision increased dramatically.

**Figure 2.**
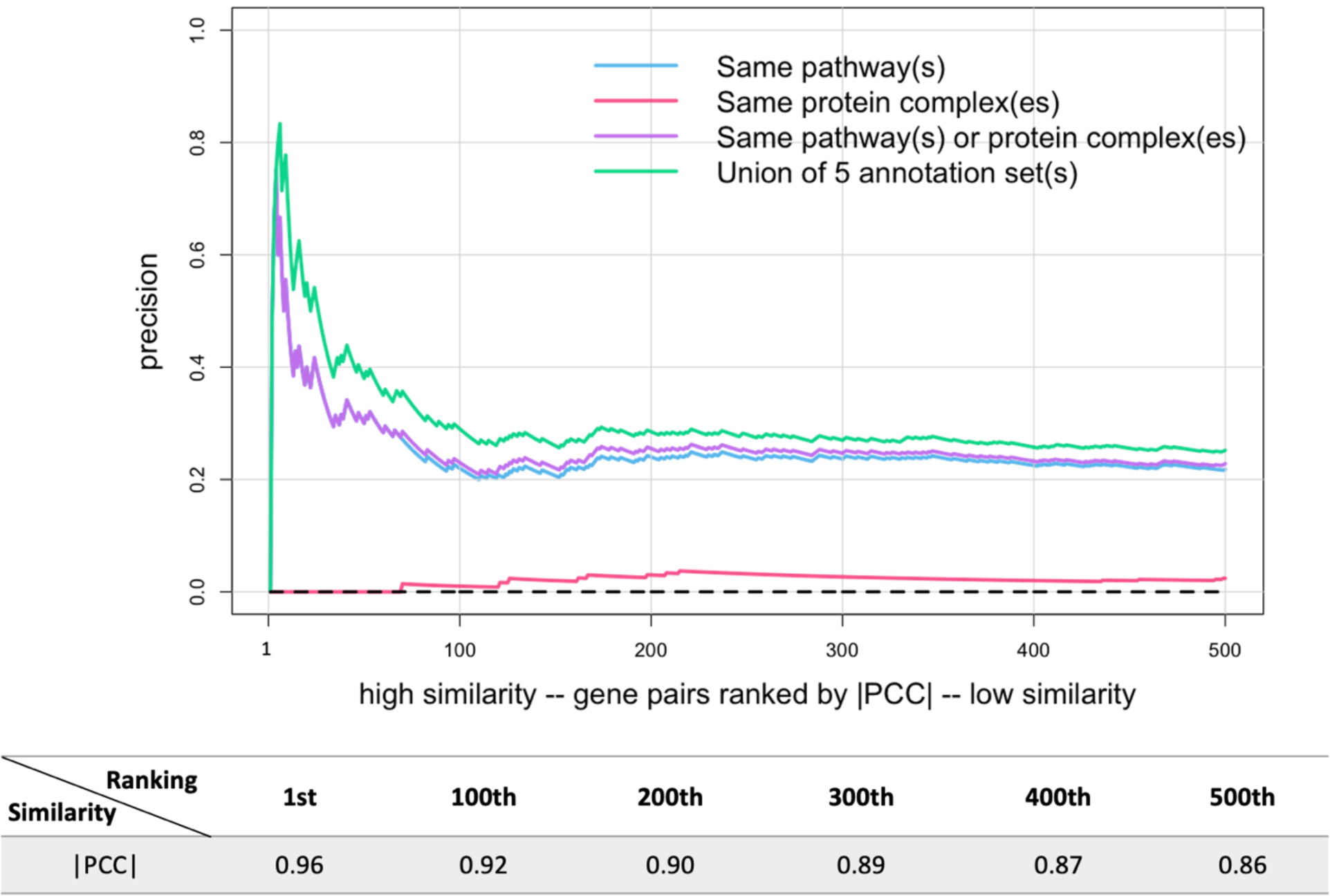
Increased co-annotation was found for gene pairs with higher phenotypic profile similarity. Gene pairs were ranked from high to low similarity based on |PCC| values and plotted versus precision [TP/(TP+FP)], which was calculated as described in the text (only the first 500 gene pairs are shown). Note that for the first few gene pairs the lines overlap except the line for protein complexes. The dashed line shows precision for randomly ordered gene pairs (negative control). The correspondence between |PCC| and ranking is shown below the graph.

### The Pearson Correlation Coefficient is sensitive to the extreme fitness scores on minimal media

To try to understand why precision was so low for protein complex annotations (Figure 2), we inspected the gene pairs and saw that 98 of the 100 top-ranking gene pairs consisted of genes coding for biosynthetic enzymes, and, in 84 of these 98 gene pairs, the genes were annotated to different biosynthetic pathways. For example, the top-ranked gene pair (|PCC| = 0.96) contained the genes *ilvC* and *argB*, which encode enzymes required for isoleucine-valine and arginine biosynthesis, respectively. Mutant strains lacking any of these biosynthetic genes would be auxotrophs and share the phenotype of little or no growth on unsupplemented minimal media. To test whether the |PCC|-based measure of phenotypic profile similarity was dominated by the large negative fitness scores associated with the auxotrophic phenotypes, we excluded the fitness scores for the growth conditions that involved minimal media (10 out of 324 total conditions) and reassessed the relationship between precision and phenotypic profile similarity. As shown in Figure 3, even though only a small fraction of conditions were excluded, this change resulted in dramatically higher precision overall, regardless of which functional annotation set was used to score co-annotation. A comparable increase in precision was also seen when auxotrophic mutants were excluded from the data set (Supplemental Figure S3).

**Figure 3.**
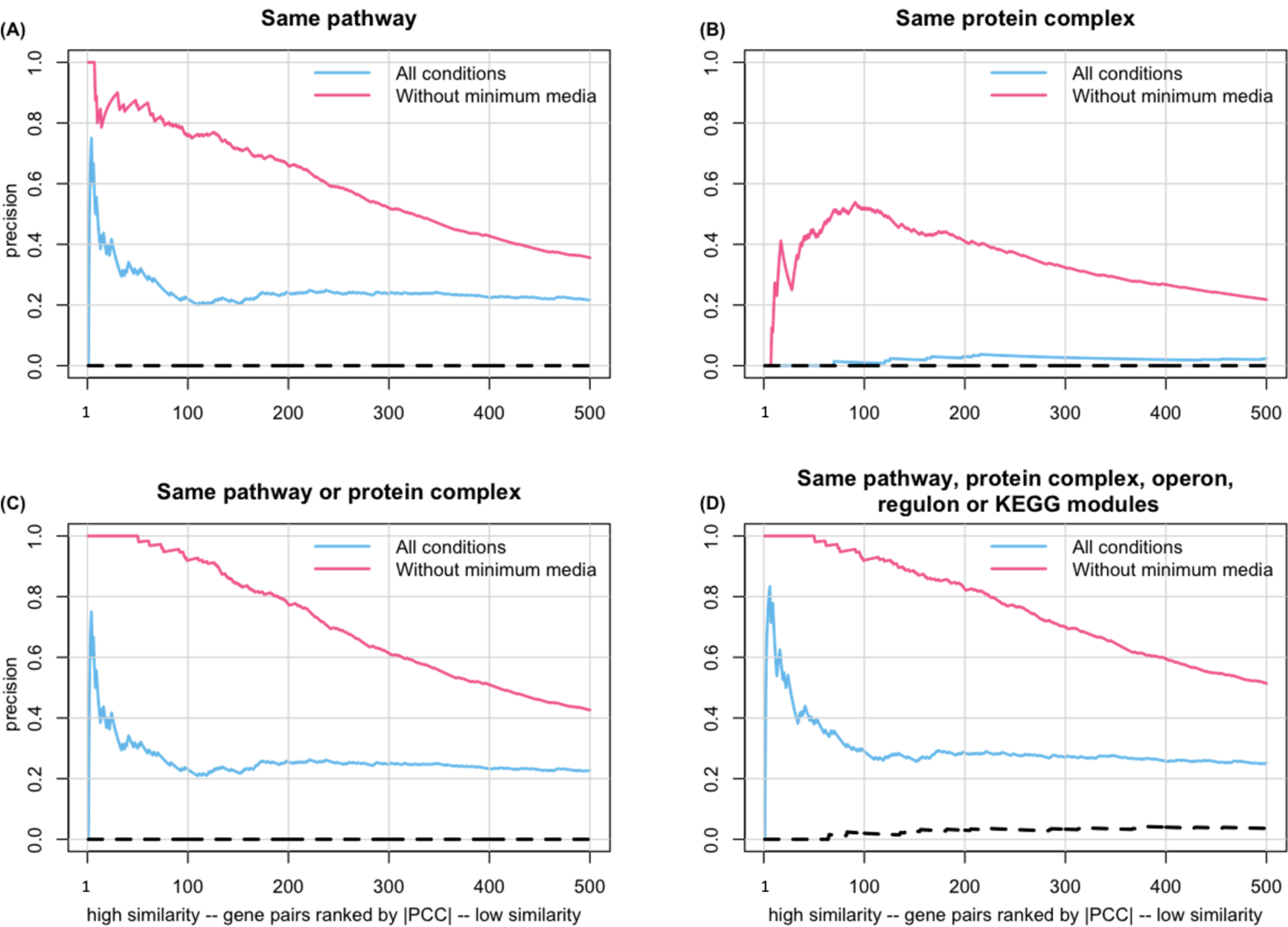
Precision increased when minimal media conditions were excluded. Gene pairs were ranked from high to low similarity based on |PCC| and plotted versus precision, calculated as described in the text (only the first 500 gene pairs are shown). The dashed line shows precision for randomly ordered gene pairs (negative control). The correspondence between |PCC| and ranking is the same as in Figure 2.

### Alternative metrics for measuring phenotypic profile similarity

There are other methods, besides the Pearson Correlation Coefficient, that can be used to assess similarity. We chose the absolute value of Spearman’s Rank Correlation Coefficient (|SRCC|) or mutual information (MI), which were implemented as described in the methods, to measure phenotypic profile similarity, and used the union of the five annotation sets to score co-annotation. Violin plots of the distributions of phenotypic profile similarity obtained using these alternative metrics were not significantly different from the distributions seen using |PCC| as the metric (results not shown). In contrast, as shown in Figure 4a, the correlation between phenotypic profile similarity and precision was dramatically higher for |SRCC| and MI compared to |PCC|. For both |SRCC| and MI, precision was >0.9 for the top 100 ranked gene pairs and remained >0.5 for approximately the top 500 pairs. This result indicates that determining phenotypic profile similarity using Spearman’s Rank Correlation Coefficient or Mutual Information is less sensitive to the presence of a relatively small number of extreme phenotype scores than using the Pearson Correlation Coefficient. If we recalculate precision for all three metrics after excluding the 10 growth conditions where auxotrophic mutants don’t grow, there is very little difference in precision for the three metrics (Figure 4b).

**Figure 4.**
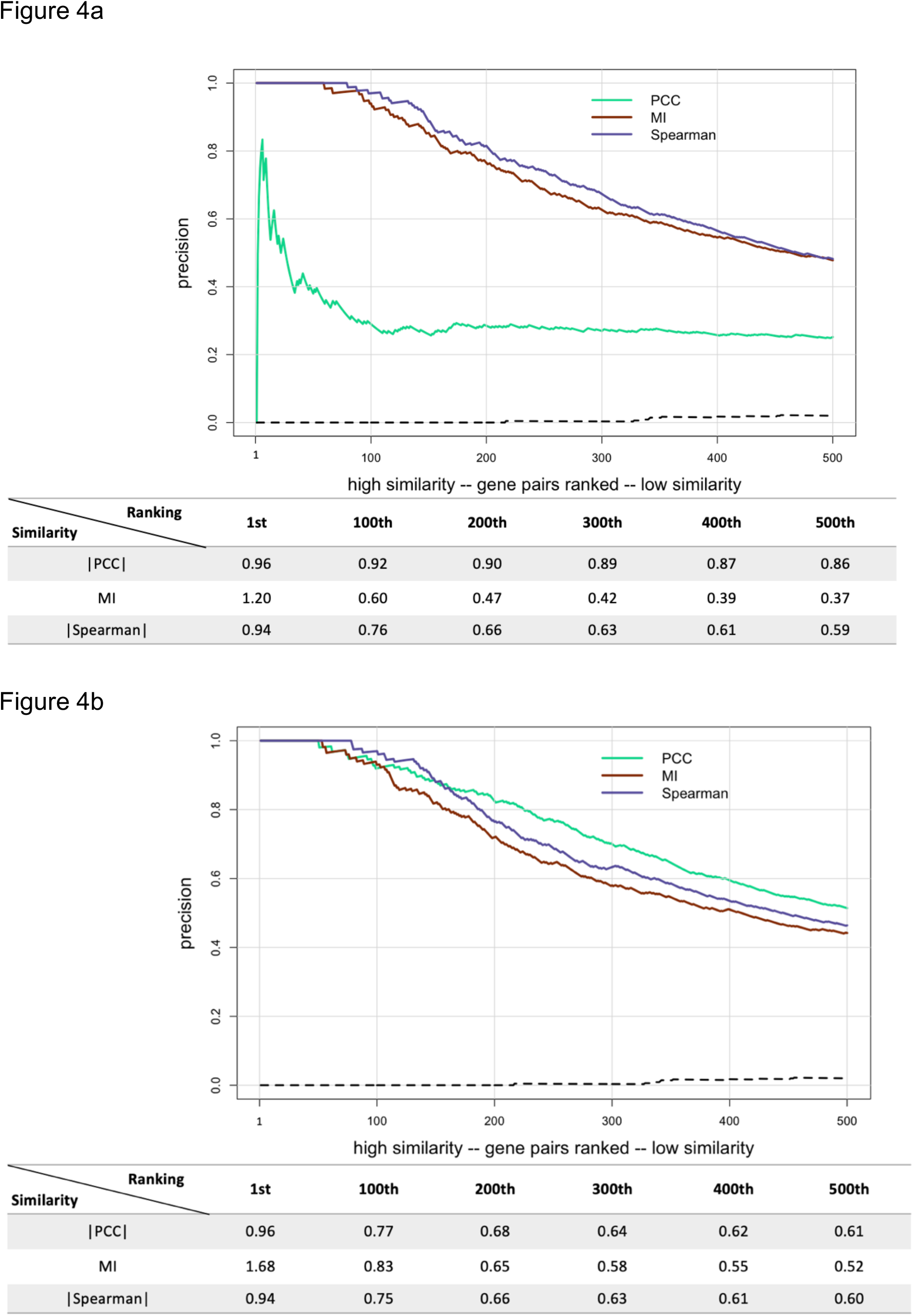
Precision versus ranking for different methods of measuring phenotype profile similarity. Gene pairs were ranked from high to low similarity and plotted versus precision, calculated as described in the text (only the first 500 gene pairs are shown). Phenotypic profile similarity was assessed using either |PCC|, MI, or |SRCC| with (a) all growth conditions used or (b) excluding growth conditions with minimal media. The dashed line shows precision for randomly ordered gene pairs (negative control). The correspondence between similarity scores and ranking is shown below each graph.

### Simplified phenotypic profiles preserve biological meanings

Combining phenotypic information from different studies is expected to increase the likelihood of finding associations between genes and functions. However, the ability to combine datasets can be limited by differences in how quantitative phenotypes are scored and by the need for methods to combine quantitative and qualitative phenotypic information. Different quantitative datasets could be combined by renormalizing the data to make them interoperable. Alternatively, quantitative phenotypes could be converted to qualitative phenotypes, which would allow integration of both quantitative and qualitative data. We chose to test the second approach because, if successful, it would allow more datasets to be combined.

The quantitative fitness scores in the phenotypic dataset were discretized to create a qualitative dataset with the fitness scores converted to 1, 0, or -1, where 1 stands for increased fitness, -1 for decreased fitness, and 0 for no difference in fitness compared to the mean fitness for all strains in a particular growth condition. The |PCC| values used to separate the three phenotype classes were based on the 5% false discovery rate as described (Nichols et al., 2011). Because the majority of strains have no significant phenotype in the growth conditions used (Nichols et al., 2011), after discretizing the data the majority of strains will have fitness scores of 0. Therefore, the Pearson Correlation Coefficient was no longer suitable for measuring phenotypic profile similarity. Instead, mutual information (MI) (Priness et al., 2007) was used as the scoring metric. The distribution of MI values for gene pairs were plotted as violin plots. The first violin plot in Figure 5a shows the distribution of MI values for all possible gene pairs, followed, from left to right, by the distribution of MI values for gene pairs co-annotated to either the same pathway; the same protein complex; the same pathway and protein complex; or the same pathway, protein complex, operon, regulon, and KEGG module. Converting the continuous quantitative fitness values to discrete ternary scores reduced the variation in the data, reflected by the change in shape of the violin plots compared to the plots shown in Figure 1. However, as was seen for the mean |PCC| values in the analysis of the quantitative data (Figure 1), the mean MI values increased as the functional associations for a given gene pair increased (Figure 5a inset).

**Figure 5.**
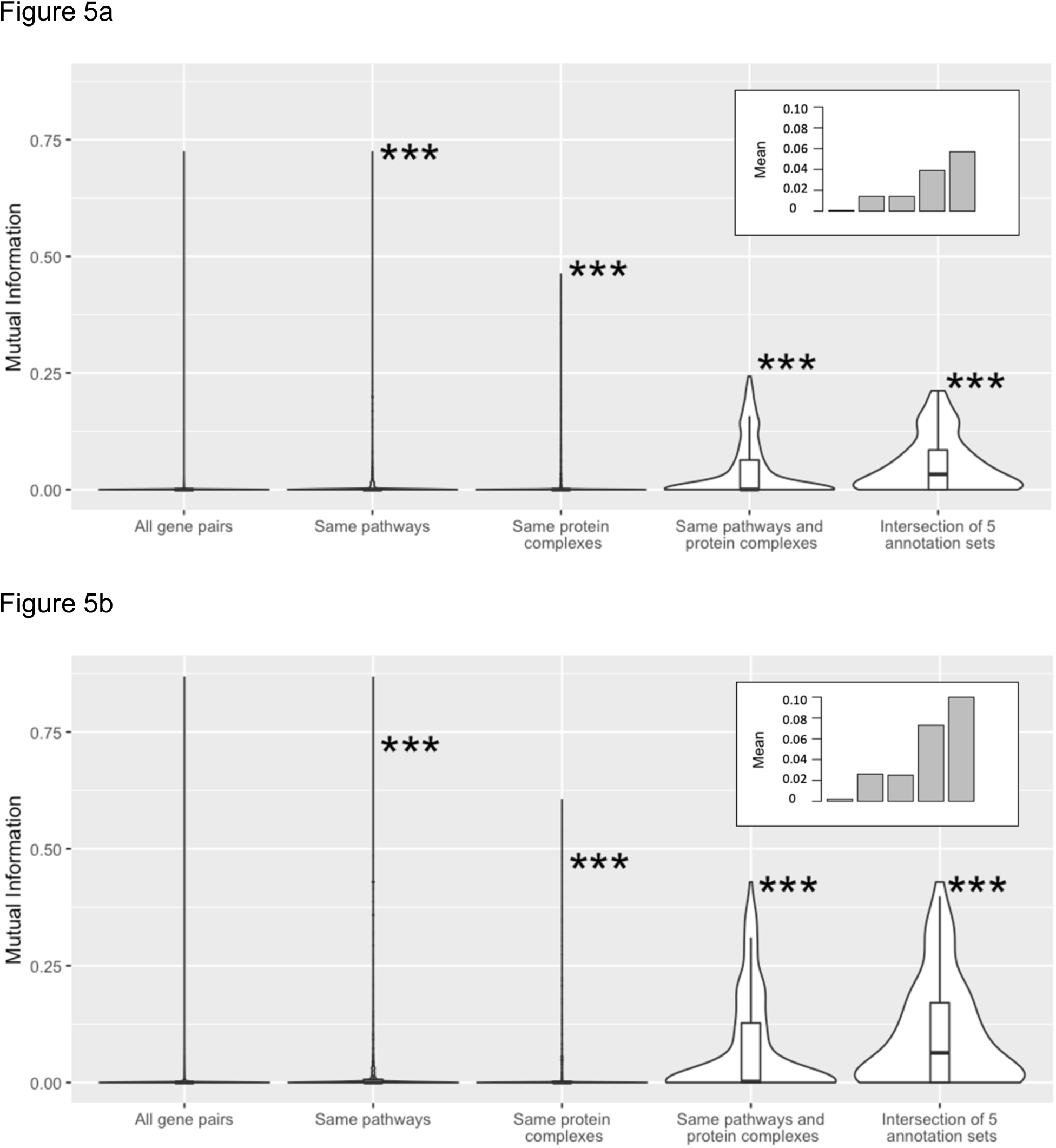
Phenotypic profile similarity after converting fitness scores from quantitative to qualitative, ternary values. Shown are violin plots of the distributions of phenotypic profile similarity based on Mutual Information for the indicated groups of gene pairs. Panel (a) shows results determined using all 324 growth conditions, and panel (b) shows results determined after collapsing the growth conditions to 114 unique stresses. The insets show the mean value for each distribution. For (a) the mean values are 0.0006, 0.014, 0.014, 0.039, and 0.057). For (b) the mean values are 0.0021, 0.026, 0.025, 0.073, and 0.1). ***: p-value <0.001 determined by 1-sided Mann-Whitney U test.

Many of the growth conditions used in the original chemical genomics study involved multiple tests of the same chemical present at different concentrations. To test the effect of further simplifying the phenotypes, the original 324 growth conditions were reduced to 114 unique stresses by including the score for only the most significant phenotype for each chemical treatment (1 or -1, as appropriate, or using a score of 0 if no significant phenotypes were seen for that treatment). The violin plots in Figure 5b show the distribution of MI values for all gene pairs and for different combinations of annotation sets for the reduced dataset. As seen for the full qualitative dataset, the mean MI values for co-annotated gene pairs in the reduced dataset were significantly higher than the mean MI value for all possible gene pairs (Figure 5b inset). In addition, when the distributions of gene pairs in the same co-annotation group are compared between Figures 5a and 5b, very significant differences of the means were observed for every co-annotated group (p-value <0.001). Overall, these results indicate that useful inferences about gene function can still be made after the conversion of quantitative phenotypes to qualitative phenotypes and even after collapsing the number of phenotypes for each chemical treatment.

We expected loss of information after quantitative phenotype scores were converted to the discretized, ternary fitness scores. To compare how many functional associations could still be retrieved using the qualitative scores, gene pairs were sorted based on their MI values determined using either quantitative phenotype scores, the qualitative ternary fitness scores, or the qualitative ternary fitness scores for the reduced set of conditions. Then precision was calculated, as described earlier, and was plotted versus ranking. As can be seen in Figure 6, precision is comparable for the top 100 gene pairs for both quantitative and discretized, qualitative fitness scores. After this point, precision drops more quickly for the qualitative data than for the quantitative data. When precision for the reduced set of conditions is compared to precision for either of the other data sets, we see that precision drops off sooner and decreases more rapidly. Yet, precision is still much higher than for randomly ordered gene pairs, which indicates that there is still significant potential in using the discretized version of phenotypes to explain functions.

**Figure 6.**
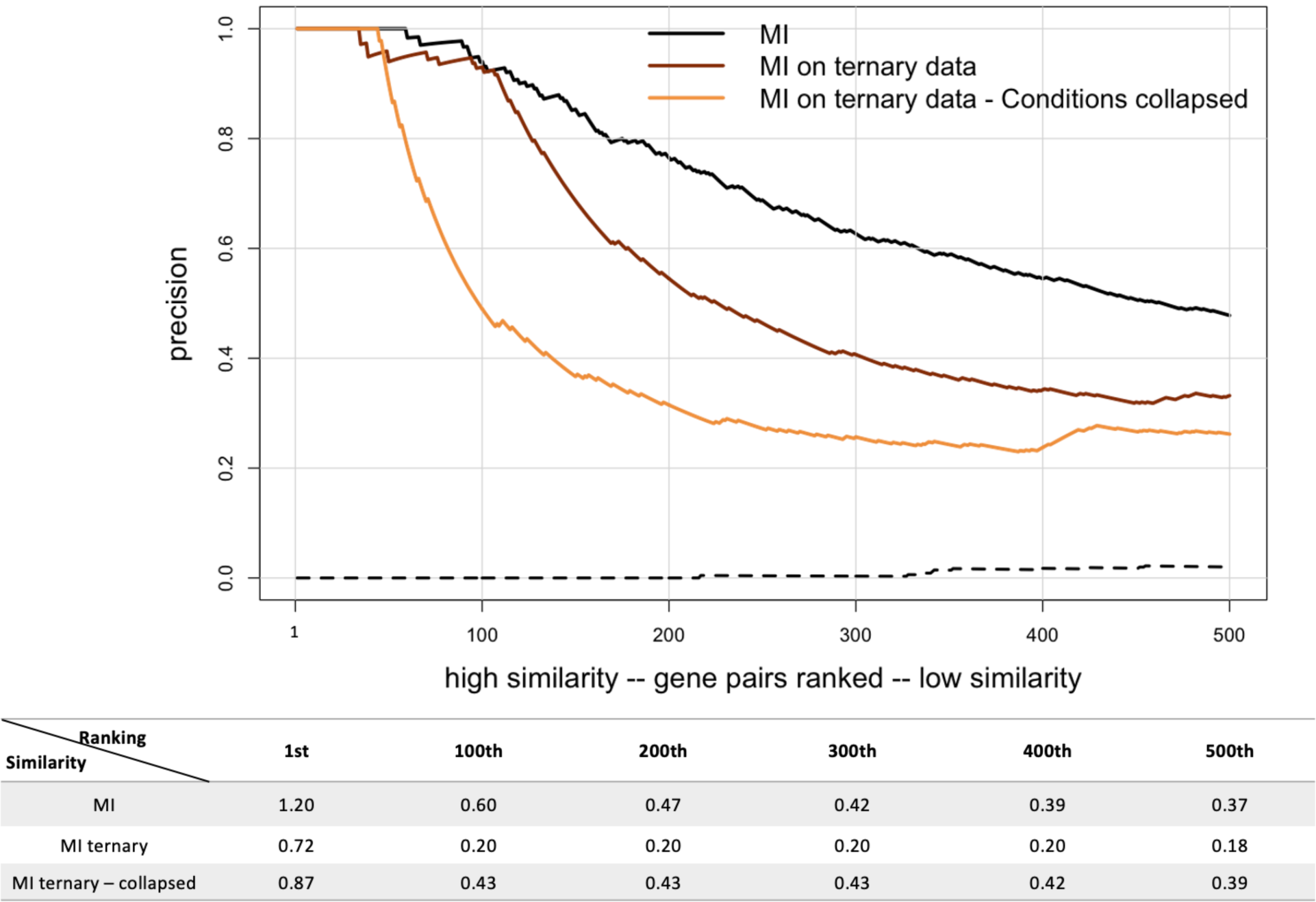
Precision versus ranking for quantitative versus qualitative, ternary fitness scores. Gene pairs were ranked from high to low similarity based on Mutual Information (MI) and plotted versus precision, calculated as described in the text (only the first 500 gene pairs are shown). The phenotypic profiles contained either the original quantitative data (black line), the discretized ternary values for all growth conditions (brown line), or the discretized, ternary values for growth conditions collapsed to 114 unique stresses (orange line). The dashed line shows precision for randomly ordered gene pairs (negative control). The correspondence between similarity scores and ranking is shown below each graph.

### Semantic similarity of GO annotations increased for gene pairs with shared functional annotations and with higher phenotypic profile similarity

Another way to assess whether two genes are likely to have similar functions is to compare the semantic similarity of the GO terms annotated to each gene. In the dataset from Nichols *et al*., 66% (2,609 out of 3,979) of the strains used have mutations of genes that are annotated with GO biological process terms, which seemed a sufficient number to justify using this approach. The Wang method (Wang et al., 2007) was used to compute semantic similarity, and the distribution of semantic similarity scores for all gene pairs where both members of the pair are annotated with at least one GO biological process term was compared to the distributions for subsets of gene pairs that have similar functions based on being co-annotated in one or more of the non-GO annotation sets. As shown in Figure 7a, semantic similarity increased when only co-annotated gene pairs were considered. The mean pairwise semantic similarity increased from 0.217 for all genes with GO biological process annotations (first violin plot) to 0.543 for gene pairs co-annotated to the same EcoCyc pathway (second violin plot), and to 0.803 for gene pairs co-annotated to the same heteromeric protein complex (third violin plot). Mean profile similarity was even higher for gene pairs that are co-annotated to both pathways and heteromeric protein complexes (mean=0.892) as well as for gene pairs that are co-annotated in all 5 annotation sets (mean=0.889), as shown in the fourth and fifth violin plots, respectively. These results show that co-annotated gene pairs are also enriched for functional similarity based on GO biological process annotations.

**Figure 7.**
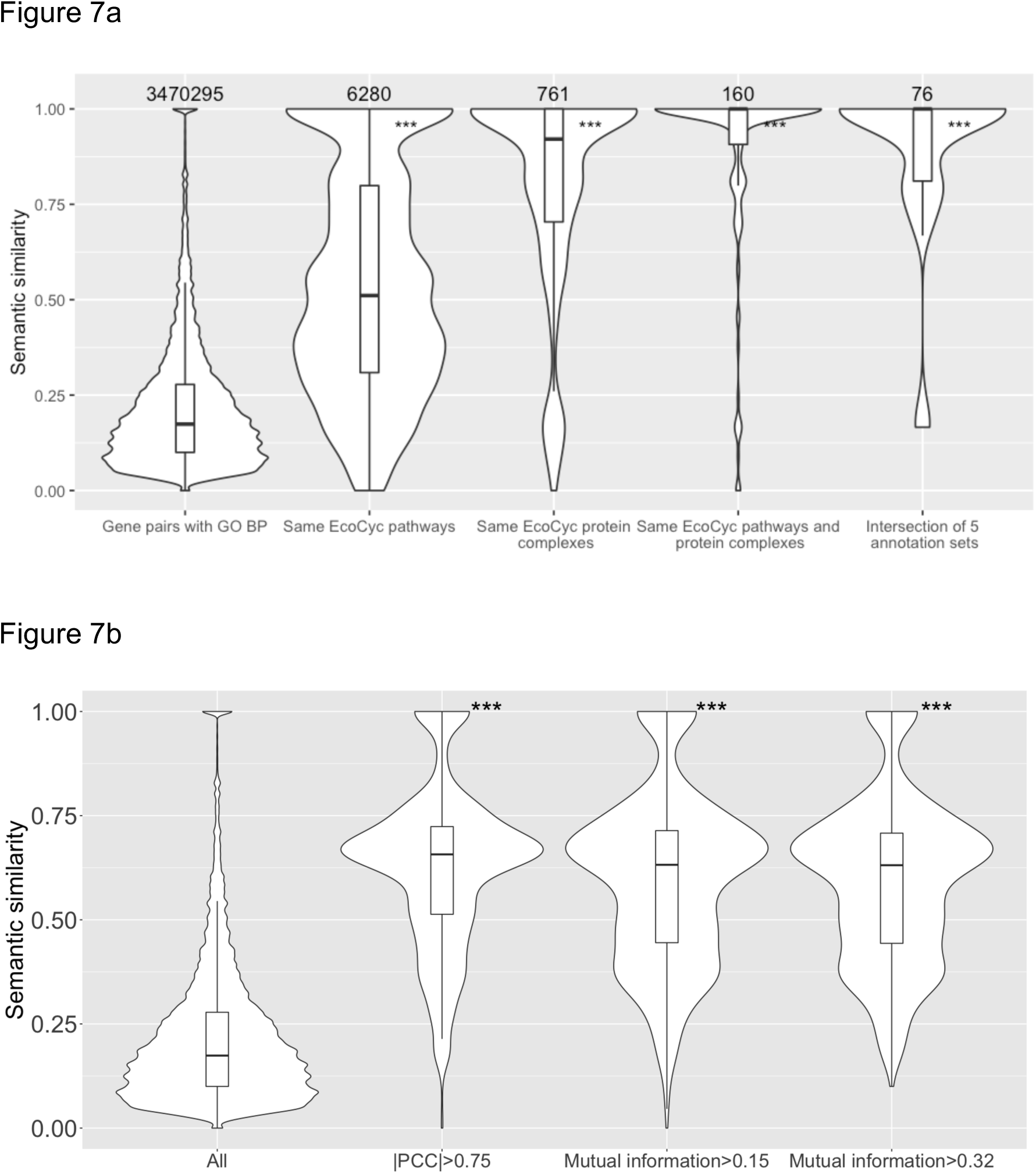
Higher semantic similarity and phenotypic profile similarity were found for co-annotated gene pairs. (a) Violin plots of the distributions of semantic similarity for the indicated groups of gene pairs. Numbers above each violin plot indicate the number of gene pairs in each plot. (b) Violin plots of semantic similarity for, from left to right: all gene pairs annotated with GO biological process term(s); the subset of gene pairs with |PCC| >0.75; the subset of gene pairs with MI >0.15 (calculated based on qualitative fitness scores for all growth conditions); and MI >0.32 (calculated based on qualitative fitness scores for the collapsed set of growth conditions). The cutoffs of MI >0.15 for the third violin plot and MI >0.32 for the fourth violin plot were chosen so that all three subsets of gene pairs would contain the same number (∼1,000) of top-ranked gene pairs. ***: p-value <0.001 was determined by 1-sided Mann-Whitney U test, compared to all gene pairs.

To test whether gene pairs that have higher phenotypic profile similarity are more likely to have similar functions based on GO biological process annotations, we compared the distributions of semantic similarity values for all gene pairs annotated with GO biological process terms and for subsets of these gene pairs that have high phenotypic profile similarity based on |PCC| or MI. The violin plots in Figure 7b show, from left to right, the distribution of semantic similarity values for all gene pairs with GO biological process annotations, the subset of gene pairs with |PCC| >0.75, the subset of gene pairs with MI >0.15 (where MI was determined using the ternary qualitative fitness scores for all growt conditions), and the subset of gene pairs with MI >0.32 (where. Comparison of the first two violin plots shows that gene pairs with |PCC| >0.75 are significantly enriched for higher semantic similarity. (The cutoff of |PCC| >0.75 was chosen arbitrarily to represent a moderate to high correlation (Hinkle et al., 2002).) Enrichment for higher semantic similarity scores was also seen for the next two subsets of gene pairs, where phenotypic profile similarity was calculated using the qualitative, ternary fitness values for either all 324 growth conditions (third violin plot) or for the collapsed set of 114 growth conditions (fourth violin plot). (The MI cutoffs of >0.15 for the third violin plot and >0.32 for the fourth violin plot were chosen so that all three subsets of gene pairs would contain the same number (∼1,000) of top-ranked gene pairs.) These results are consistent with those in Figure 4b, which show higher phenotypic profile similarity enriches for co-annotated gene pairs.

In order to assess whether gene pairs that have higher semantic similarity also have higher phenotypic profile similarity, we chose an arbitrary cutoff of 0.5 for semantic similarity and used it to select a subset of gene pairs from the entire set of gene pairs with GO biological process annotations. We then compared the distribution of semantic similarity scores for the two sets of gene pairs. The violin plots are shown in Figure 8. Although the two distributions appeared almost identical, the subset of gene pairs with semantic similarity >0.5 is enriched for gene pairs with higher phenotypic profile similarity. The difference in the mean |PCC| values for the two distributions is small (0.093 vs 0.10), but it is statistically significant based on the Mann-Whitney test, p<0.0001. This is consistent with Figure 1, where co-annotated gene pairs show enriched phenotypic similarity.

**Figure 8.**
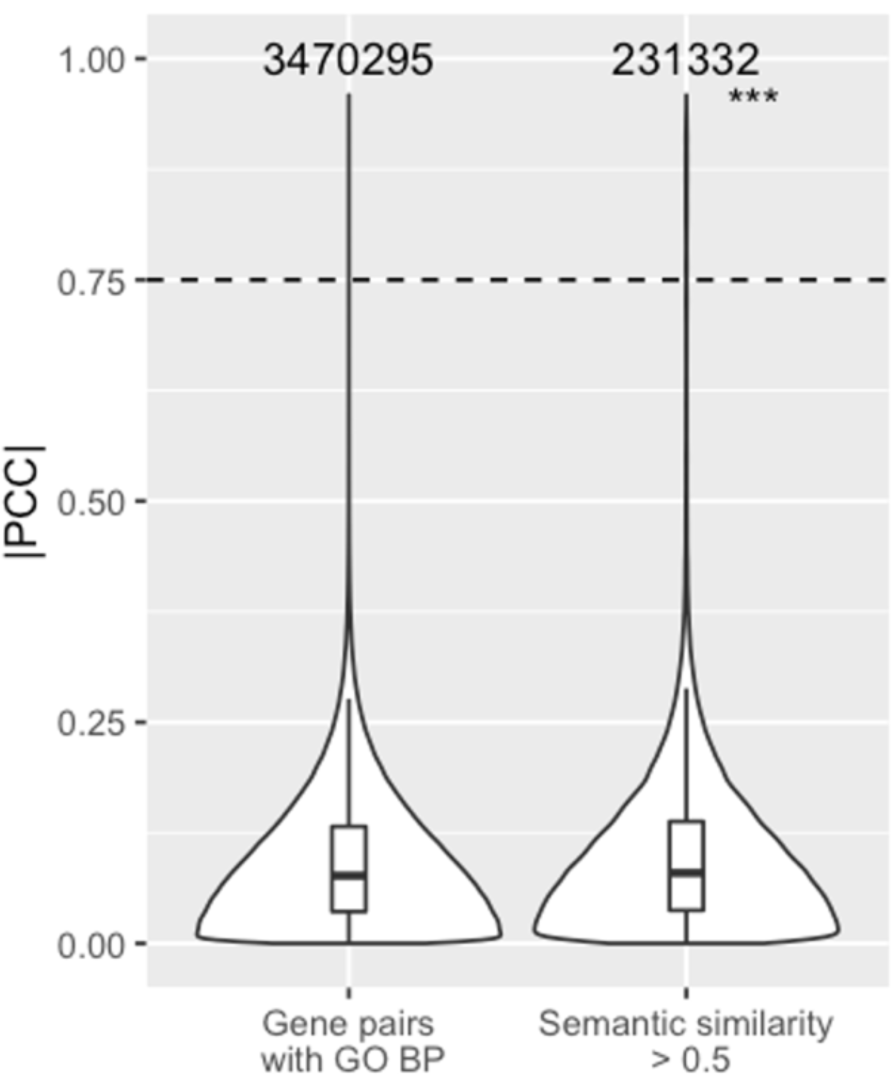
Higher phenotypic similarity was found for gene pairs that have higher GO semantic similarity. Violin plots of the distributions of the |PCC| values for all gene pairs with GO biological process annotations and the subset with semantic similarity is greater than an arbitrary cutoff of 0.5. Numbers above each violin plot indicate the number of gene pairs in each plot. ***: p-value <0.001 was determined by 1-sided Mann-Whitney U test, compared to all gene pairs.

## DISCUSSION

We systematically reanalyzed a published high-throughput phenotypic profile dataset for the model Gram-negative bacterium *E. coli* comparing different metrics for measuring phenotypic profile similarity, and assessing the effect of converting quantitative fitness scores to qualitative fitness on measurements of phenotypic profile similarity. We re-examined the *E. coli* phenotypic profiles in a pairwise fashion with the help of existing functional annotations. Overall, we found that gene pairs with functional associatons are enriched for high phenotypic profile similarity scores and that gene pairs with high phenotypic similarity scores tend to have functional associations.

Six high-quality annotations sets were used as sources of functional information. The gene annotations in EcoCyc, RegulonDB, KEGG, and GO come primarily from expert manual curation (Gama-Castro et al. 2016; Kanehisa et al. 2016; Keseler et al. 2017; Keseler, 2014; Gene Ontology Consortium, 2017). The GO biological process annotations include ∼1,200 annotations (21%) that are inferred from electronic annotation without additional human review. We decided to include the electronic annotations in our analysis because most of them come from the transfer of annotations from orthologous gene products or are based on mappings from external sources, such as InterPro2GO or EC2GO, which have been shown to be very accurate (Camon et al. 2005; Hill et al. 2001; Holliday et al. 2017). Indeed, there was no significant difference in the semantic similarity of gene pairs whether electronic annotations were included (Figure 7b) or excluded (Figure S4).

One aim of this study was to determine whether different metrics for determining phenotypic profile similarity differed in their ability to identify gene pairs with functional similarity. We compared the performance of the metrics based on precision: the fraction of positive results that are true positives. Gene pairs with phenotypic profile similarity above a specified cutoff were considered as positive results, and true positives were defined as gene pairs that are co-annotated in at least one of the five annotation sets. We chose to use precision rather than accuracy, which is the fraction of correct results, because the co-annotated and non-co-annotated gene pairs constitute a highly imbalanced dataset (Saito & Rehmsmeier, 2015). Because the number of non-co-annotated gene pairs is much larger than the number of co-annotated gene pairs, high accuracy could be achieved by classifying all gene pairs as true negatives, but this wouldn’t be very informative.

Overall, there appeared to be little difference in the performance of |PCC|, |SRCC| or MI based on their precision scores for the top 500 gene pairs (Figure 4b). Initially, it appeared that |SRCC| and MI outperformed |PCC| (Figure 4a). However, when the analysis was repeated after removing the conditions involving growth on minimal media, the precision for gene pairs ranked based on |PCC| increased significantly (compare Figures 4a and 4b). We suggest that this difference is due to the sensitivity of the Pearson Correlation Coefficient to outliers in the data (Schober et al., 2018). We realized that the collection of strains used by Nichols et al. contains many mutants that have little or no growth on minimal media because the gene for a biosynthetic enzyme is deleted. In contrast, these auxotrophic mutants didn’t have a significant phenotype in most of the other growth conditions tested, which used rich media, so the large negative fitness scores on minimal media were essentially outliers. In our analysis, the sensitivity of PCC to outliers interfered with the measurement of precision because there were so many combinations of genes from different biosynthetic pathways that shared an auxotrophic phenotype but did not share a functional annotation in the annotation sets used.

However, this doesn’t mean that |PCC| can’t be used to measure phenotypic profile similarity in high-throughput phenotype screens. For most gene pairs that don’t include an auxotrophic mutant, the phenotypic profile similarity (based on |PCC|) changed very little when minimal media conditions were removed (data not shown). However, there were a few gene pairs where a possible functional association could have been missed if the minimal media conditions were not removed. We illustrate this with a gene pair where the functions of the gene products are known to have a functional association. The *exbD* and *fepA* genes are both needed for transport of ferric iron-enterobactin across the outer membrane (Noinaj et al. 2010). When profile similarity was calculated using the fitness scores for all conditions, |PCC| = 0.4773. After minimal media conditions were removed, |PCC| increased to 0.6204, a high enough correlation that this gene pair would be a reasonable candidate for future experiments.

In addition to showing comparable precision, the three metrics, |PCC|, |SRCC|, and MI, also produced comparable profile similarity scores for many, although not all, gene pairs. We conclude that there is no single best way to measure phenotypic profile similarity. Instead, it may be advantageous to use more than one correlation metric when searching for functional associations. For high-throughput experiments that measured growth of a large number of strains in many different environments, it may also be useful to preprocess the fitness data, such as filtering or combining results from certain growth conditions.

To make it easier to compare results for the different similarity metrics, we have made the data set from Nichols et al. available in a searchable, interactive format that allows queries for strains, conditions, and phenotypic profile similarity of gene pairs determined by |PCC|, |SRCC|, MI, and semantic similarity (https://microbialphenotypes.org/wiki/index.php?title=Special:Ecolispecialpage).

The relationship between precision and ranking based on profile similarity shown in Figure 4b suggests that a shared function is known for most of the highly correlated gene pairs. To test this idea, we used a cutoff of |PCC| >0.75 to define highly correlated gene pairs, filtered out the gene pairs that have no co-annotations, and then manually examined the gene pairs. If fitness scores for the growth conditions involving minimal media were excluded, there were only 10 non-co-annotated gene pairs (summarized in Table 1). We found functional associations that could explain the observed phenotypic profile similarity for 7 of the 10 gene pairs. In one case, the two genes (*dsbB* and *dsbA*) showed up as non-co-annotated because they are in a pathway that wasn’t yet included in EcoCyc version 21.1. The other six gene pairs highlight some of the challenges of creating (and using) annotation, such as deciding where pathways start and end and determining appropriate levels of granularity. For example, the gene pairs *rfaF*(*waaF*)-*rfaE*(*hldE*) and *rfaF*(*waaF*)-*lpcA* (*gmhA*) are non-co-annotated, even though all three genes are required for synthesis of the lipid A-core oligosaccharide component of outer membrane lipopolysaccharide. The explanation is that *rfaF*(*waaF*) is annotated to the central assembly pathway for building the lipid-core oligosaccharide moiety, while *rfaE*(*hldE*) and *lpcA*(*gmhA*) are annotated to a branch pathway that builds one of the saccharide subunits of the core (Raetz & Whitfield, 2002). The functional association between the three genes would have been revealed if we had included GO annotations, since all three genes are annotated to the GO term for the lipopolysaccharide core region biosynthetic process (GO:0009244).

**Table 1.** Non-co-annotated gene pairs with |PCC| >0.75

| Gene pair<sup>1</sup> | Known or predicted functional association |
| --- | --- |
| ECK0730-*pal*_ECK0725-*ybgC*<sup>2</sup> | Tol-Pal cell envelope complex (CPLX0-2201) |
| ECK0768-*uvrB*_ECK2563-*recO* | DNA repair (recombinational repair RECFOR-CPLX and nucleotide excision repair UVRABC-CPLX) |
| ECK1912-*uvrC*_ECK2563-*recO* | DNA repair (recombinational repair RECFOR-CPLX and nucleotide excision repair UVRABC-CPLX) |
| ECK2901-*visC(ubil)*_ECK3033-*yqiC(ubiK)*<sup>3</sup> | ubiquinol-8 biosynthesis (PWY-6708) |
| ECK3610-*rfaF(waaF)*_ECK3042-*rfaE(hldE)*<sup>4</sup> | superpathway of lipopolysaccharide biosynthesis (LPSSYN-PWY) |
| ECK3610-*rfaF(waaF)*_ECK0223-*lpcA*<sup>4</sup> | super pathway of lipopolysaccharide biosynthesis (LPSSYN-PWY) |
| ECK3852-*dsbA*_ECK1173-*dsbB* | periplasmic disulfide bond formation (PWY0-1599)<sup>5</sup> |
| ECK1544-*gnsB*_ECK2394-*gltX* | unknown |
| ECK2066-*yegK(pphC)*_ECK0345-*mhpB* | unknown |
| ECK3699-*mnmE*_ECK0050-*apaH* | unknown |

<sup>1</sup> The strain names are from supplemental Table S2 of Nichols et al. (2011). Where the gene name has changed, the new gene name is included in parentheses.
<sup>2</sup> *ybgC* is in an operon that also includes the genes for three of the protein components of the Tol-Pal cell envelope complex
<sup>3</sup> *ubiK* codes for an accessory protein required for efficient synthesis of ubiquinol-8 under aerobic conditions, but is not annotated as part the ubiquinol-8 biosynthesis pathway
<sup>4</sup> *rfaE(hldE)* and *lpcA* are not annotated to the super pathway of lipopolysaccharide biosynthesis (LPSSYN-PWY)
<sup>5</sup> PWY0-1599 was not present in EcoCyc version 21.1

We did not find a shared function for the last three non-coannotated gene pairs. Given that so many of the other highly correlated gene pairs do share a function, it is possible that future experiments will uncover a shared function for these three gene pairs. However, it also possible that the observed phenotypic profile similarity is fortuitous, as we saw for mutants with an auxotrophic phenotype or mutants with increased sensitivity to DNA damage. For example, this may be the most likely explanation for the phenotypic similarity of the *mnmE* and *apaH* genes. Both are required for growth at pH 4.5 (Nichols et al. 2011, Vivijs et al., 2016), but appear to function independently. MnmE, partnered with MnmG, modifies 2-thiouridine residues in the wobble position of tRNA anticodons (Elseviers et al., 1984), while ApaH is a diadenosine tetraphosphatase (Guranowski et al., 1983) and mRNA decapping enzyme (Luciano et al., 2019). Both MnmE and ApaH are proposed to affect resistance to pH and other stresses through their effects on gene expression (Dedon & Begley, 2014, Vivijs et al., 2016, Luciano et al., 2019).

A significant conclusion from this study is that functional associations can still be inferred from phenotypic profiles after quantitative fitness scores are converted to qualitative, ternary fitness values. While some information was lost compared to using quantitative fitness scores, the precision based on qualitative fitness values was much greater than for randomly ordered gene pairs (Figure 6). This result suggests that inherently qualitative phenotypes, such as aspects of cell morphology, could be incorporated into phenotypic profiles and used to infer functional associations. It may also be possible to incorporate phenotype annotations into phenotypic profiles. These annotations typically capture information in a qualitative fashion and have previously been shown to be useful for inferring gene function (Hoehndorf et al., 2013; Ascensao et al., 2014). These results also suggest that using qualitative phenotypes may be a viable option for integrating phenotype information from different studies. Thus, we believe that using qualitative phenotypes to combine more *E. coli* datasets, or datasets from other microorganisms, will allow us to extract many more functional insights.

## MATERIALS & METHODS

### Sources of data

The high-throughput phenotypic profiling data as normalized fitness scores were downloaded from supplemental Table S2 of the original paper (Nichols et al., 2011). Missing values (0.17% of total fitness scores) were replaced with population mean as an imputation method.

Six annotation sets including GO annotations were obtained from various sources: From a downloaded version of EcoCyc version 21.1 (http://bioinformatics.ai.sri.com/ecocyc/dist/flatfiles-52983746/), the ECK identifiers in supplemental Table S2 from the original research paper (Nichols et al., 2011) were verified, corrected and mapped to EcoCyc gene identifiers and b numbers using information in the file genes.txt. EcoCyc Pathway annotations were mapped to each gene using information in the file pathways.col. EcoCyc Protein complex annotations were mapped to each gene using information in the file protcplxs.col. KEGG module annotations were obtained and mapped by retrieving module name and b numbers from the KEGG website (https://www.kegg.jp). Operon and regulon annotations were obtained and mapped to each gene using a download of Regulon DB version 9.4 (http://regulondb.ccg.unam.mx). The file operon.txt was the source of operon annotations. The file object_synonym.txt was used to map ECK12 gene identifiers to ECK gene identifiers. RegulonDB annotations were then obtained from the file regulon_d_tmp.txt and mapped to ECK identifiers. GO biological process annotations were obtained from the Ecocyc file gene_association.ecocyc and mapped to each gene to produce the file 2017_05_ECgene_association.ecocyc.csv. UniProt IDs retrieved from the Bioconductor package UniProt.ws were used to associate GO annotations from proteins to genes. The number of genes annotated by each annotation set and the total number of annotations are shown in Table 2.

**Table 2.** Annotation sets used in this study

| Annotation set (source) | Subset of annotated genes tested<sup>a</sup> | Total no. of annotations for each subset<sup>b</sup> |
| --- | --- | --- |
| Pathways (EcoCyc) | 885 | 2,317 |
| Heterooligomeric protein complexes (EcoCyc) | 688<sup>c</sup> | 871<sup>c</sup> |
| Operons (RegulonDB) | 3,858 | 5,349 |
| Regulons (RegulonDB) | 1,572 | 3,886 |
| Modules (KEGG) | 333 | 524 |
| Pathways or Protein complexes | 1,385 | 3,269 |
| Pathways and Protein Complexes | 188 | 818<sup>d</sup> |
| Any (Union of all 5 annotation sets) | 3,866 | 12,937 |
| All (Intersection of all 5 annotation sets) | 77 | 922<sup>d</sup> |
| GO biological process | 2,609 | 5,775 |
| <sup>a</sup> Number of annotated genes that were deleted or otherwise mutated in the Nichols strain set (Nichols et al., 2011).<br><sup>b</sup> Total number of annotations associated with the genes in the first column.<br><sup>c</sup> This excludes 681 genes annotated to protein complexes whose products form only homooligomeric complexes<br><sup>d</sup> This is the number of annotations associated with any of the 77 genes that are annotated to all 5 annotation sets. |  |  |

### Statistical analysis and software

The statistical programming language R was used throughout the study. Phenotypic profile similarity was calculated using Pearson Correlation Coefficient (|PCC|), Spearman’s Rank Correlation Coefficient (|SRCC|), Mutual Information, and semantic similarity. Pearson and Spearman’s Rank Correlation Coefficients were calculated using the cor() function, with the metric argument specified by either “pearson” or “spearman”. Different implementations are needed to calculate Mutual Information for continuous, quantitative data and discretized, qualitative data. Mutual Information for quantitative data was calculated using the cminjk() function provided in the mpmi package, while Mutual Information for discretized data was calculated using the mutinformation() function provided in the infotheo package. Both packages are available from CRAN (https://cran.r-project.org/web/packages/mpmi/index.html). The semantic similarity of GO biological process annotations was calculated using a graph-based method (Wang et al., 2007). Calculations were performed using the GOSemSim package (Yu et al., 2010) from Bioconductor. For the Mann-Whitney U test, wilcox.test() function was used. For violin plots, geom_violin() was used to plot the kernel density plot and geom_box() was used for the boxplot. Both functions are from the ggplot2 package (Wickham, 2016). In the box plots associated with each violin plot, the middle lines in the boxes represent medians; the whiskers indicate the 1.5 interquartile range (IQR) away from either Q1 (lower box boundary) or Q3 (upper box boundary).

The code and data files used for calculations and reproducing the results are available on GitHub: https://github.com/peterwu19881230/Systematic-analyses-ecoli-phenotypes.

## ACKNOWLEDGEMENTS

We mourn the unexpected death of JCH who led this project and passed away on 23 Jan 2020. We hope this publication will continue his scientific legacy. We thank Michelle Giglio, Matthew Sachs and Yen-Ting Lu for helpful comments on the manuscript. This work was supported by a grant from the National Institutes of Health (R01GM089636) to JCH.

## AUTHOR CONTRIBUTIONS

JH and DS conceptualized the project. JH, PW, and DS designed the experiments and the analytical pipeline. PW implemented the experiments and analyzed the data. CR helped with the implementation of experiments. PW, DS, and JH wrote the manuscript.

## Supplemental Tables and Figures

**Supplemental Table S1.**
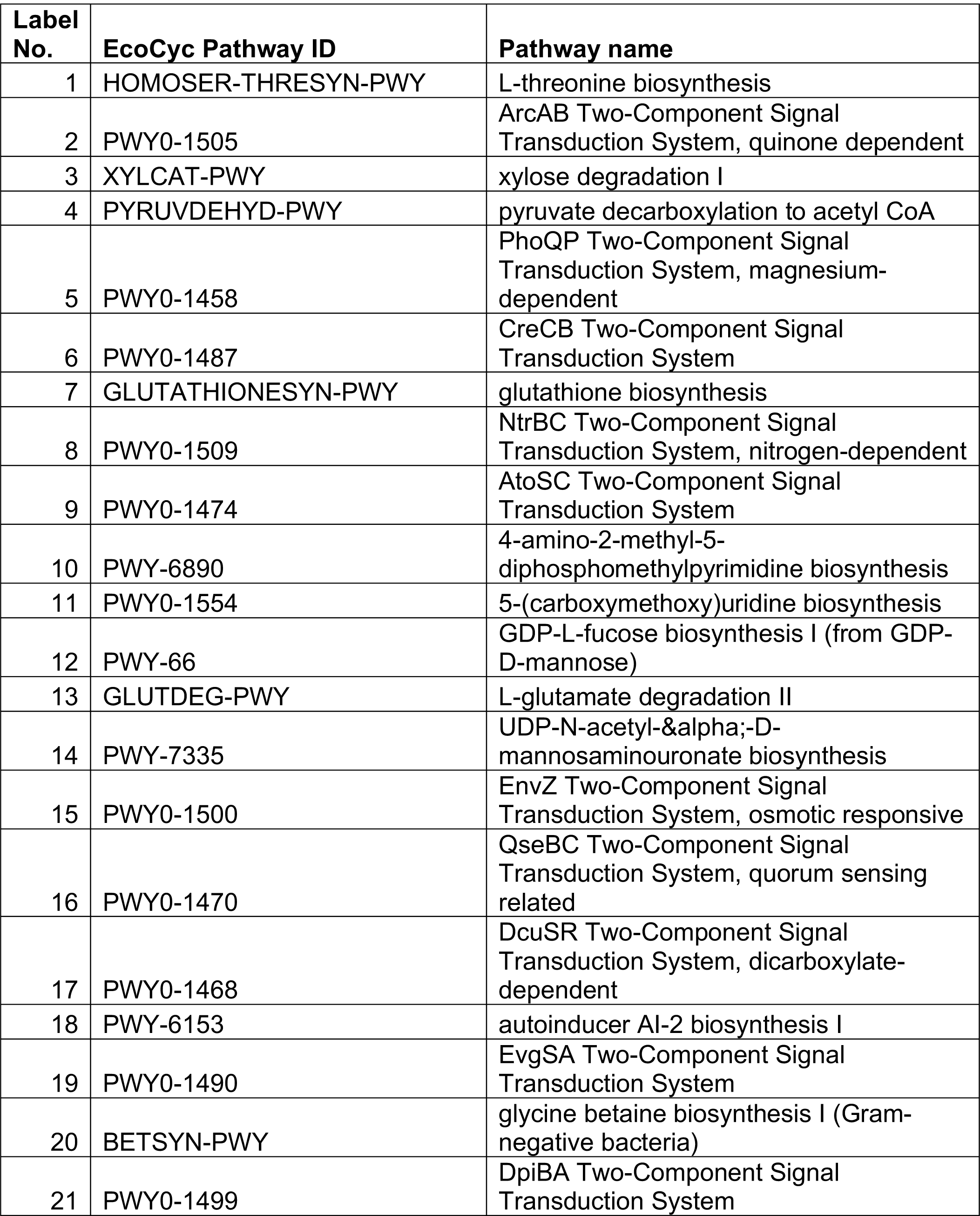
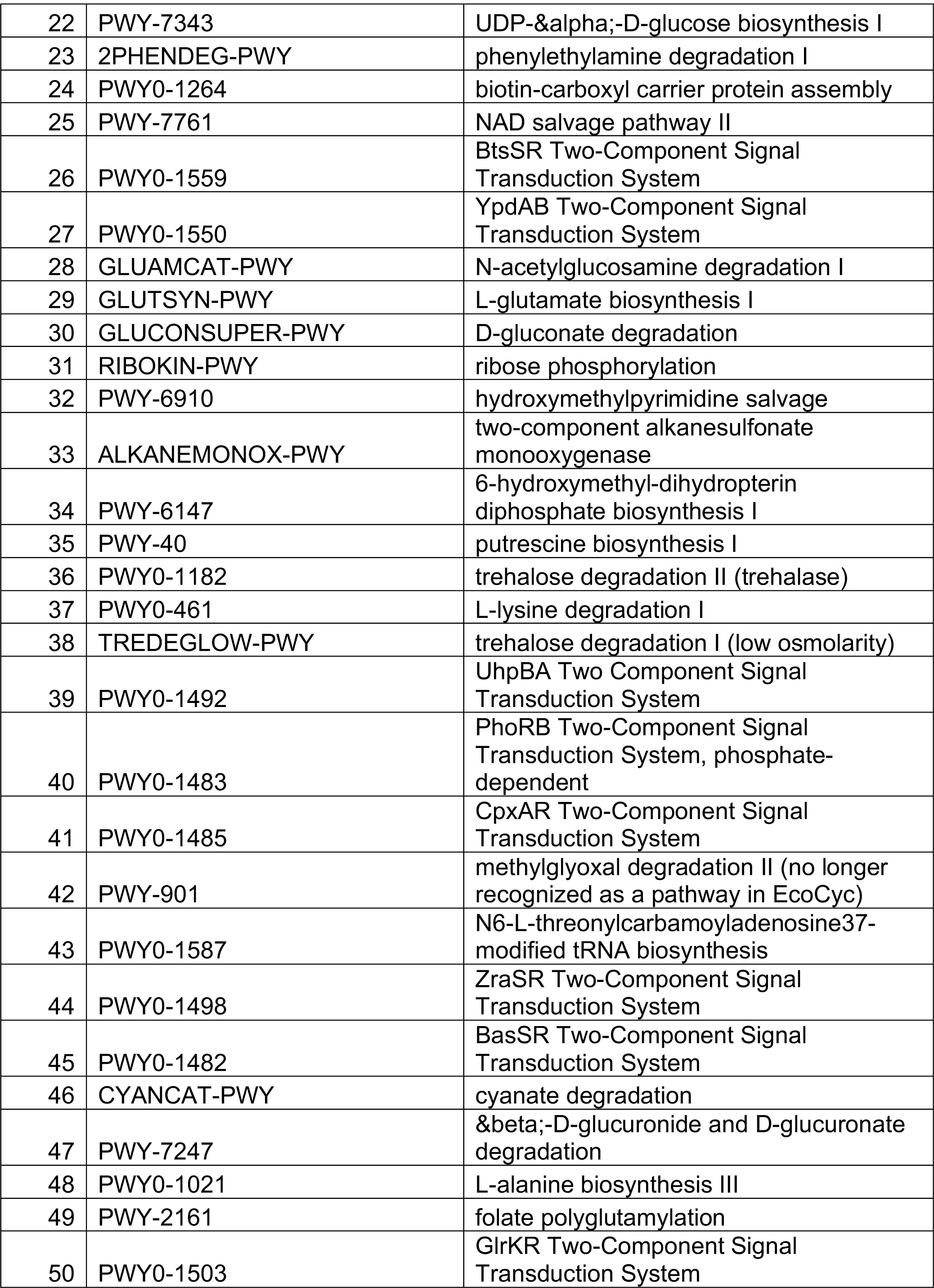
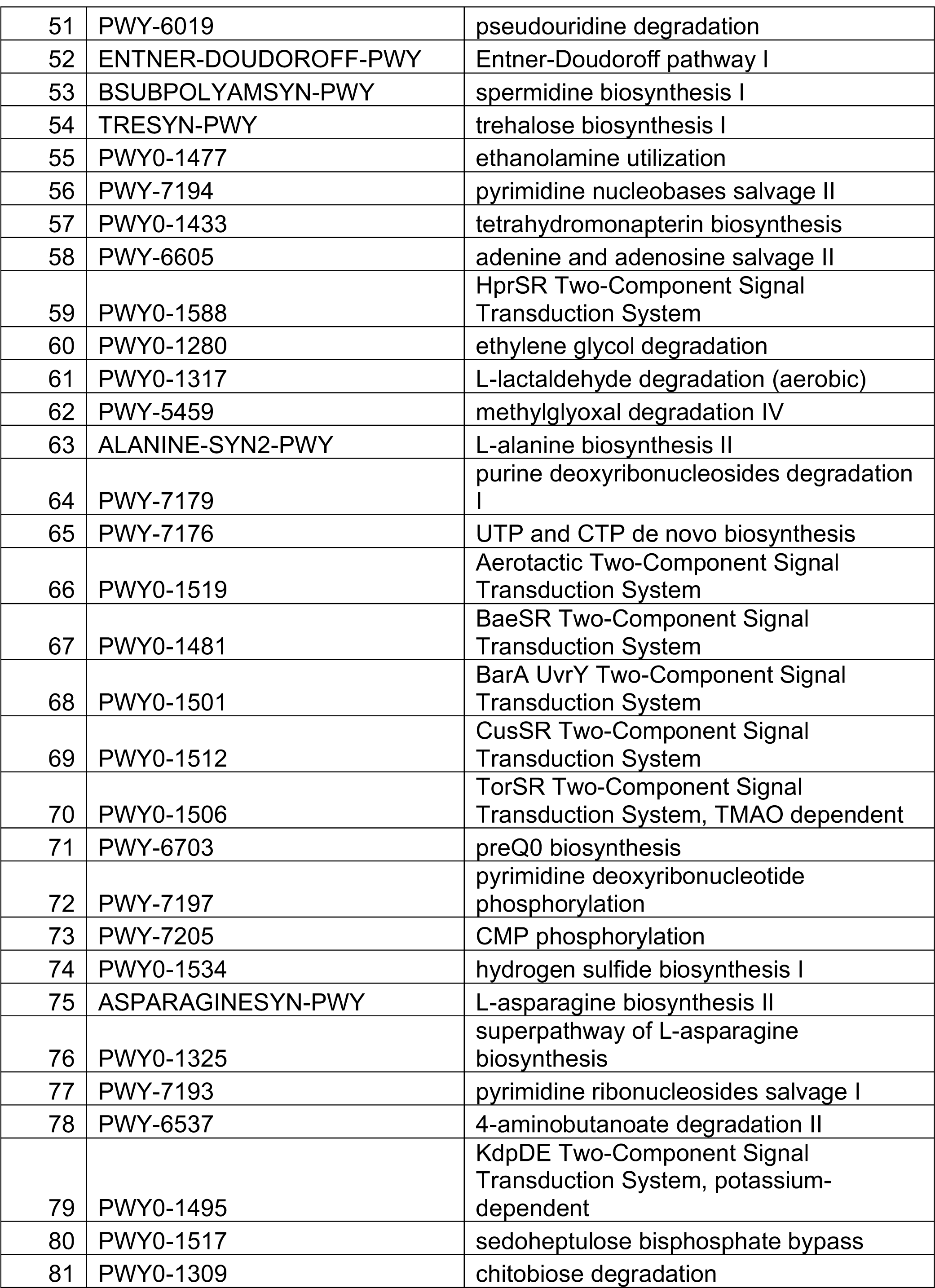
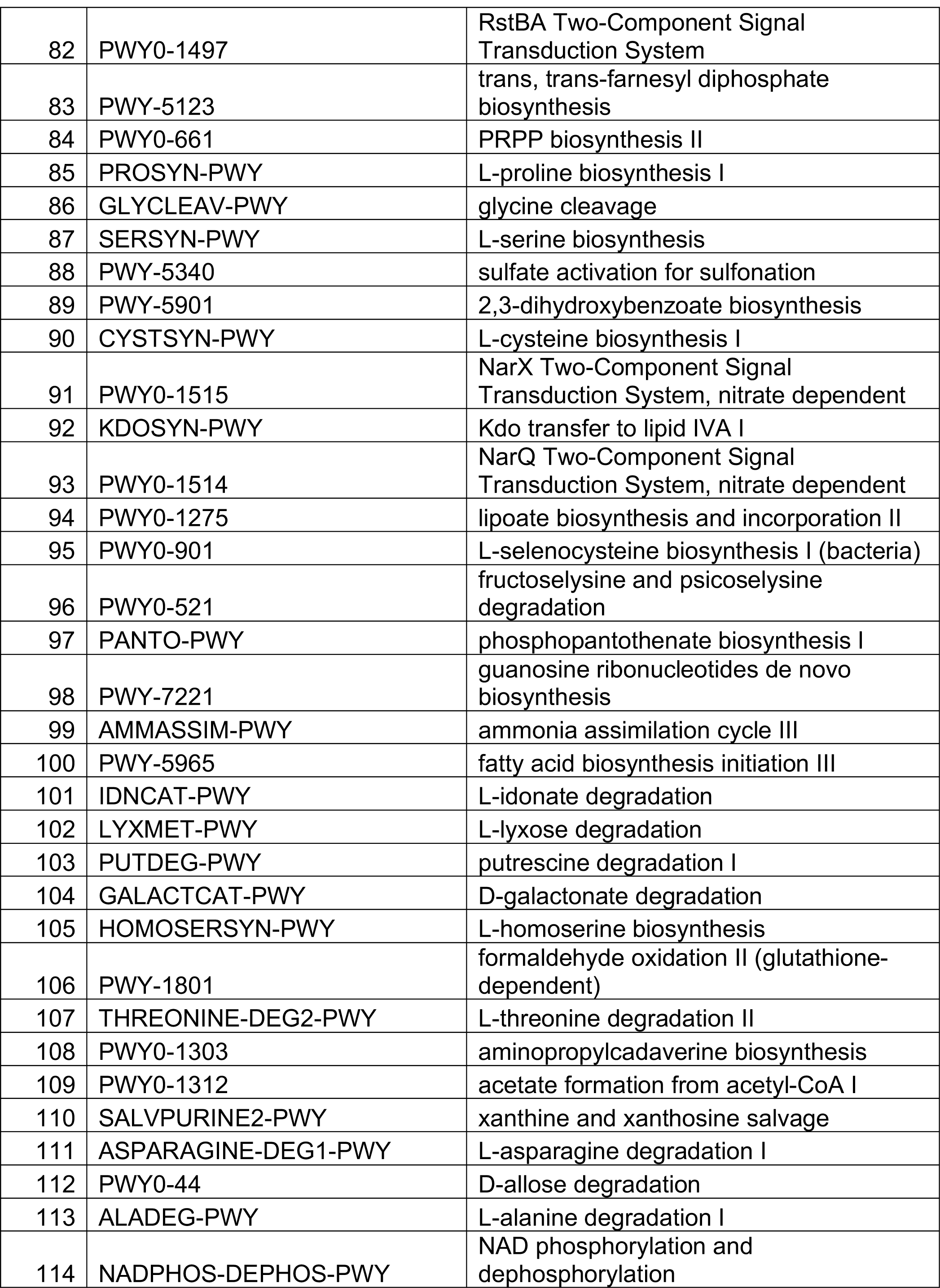
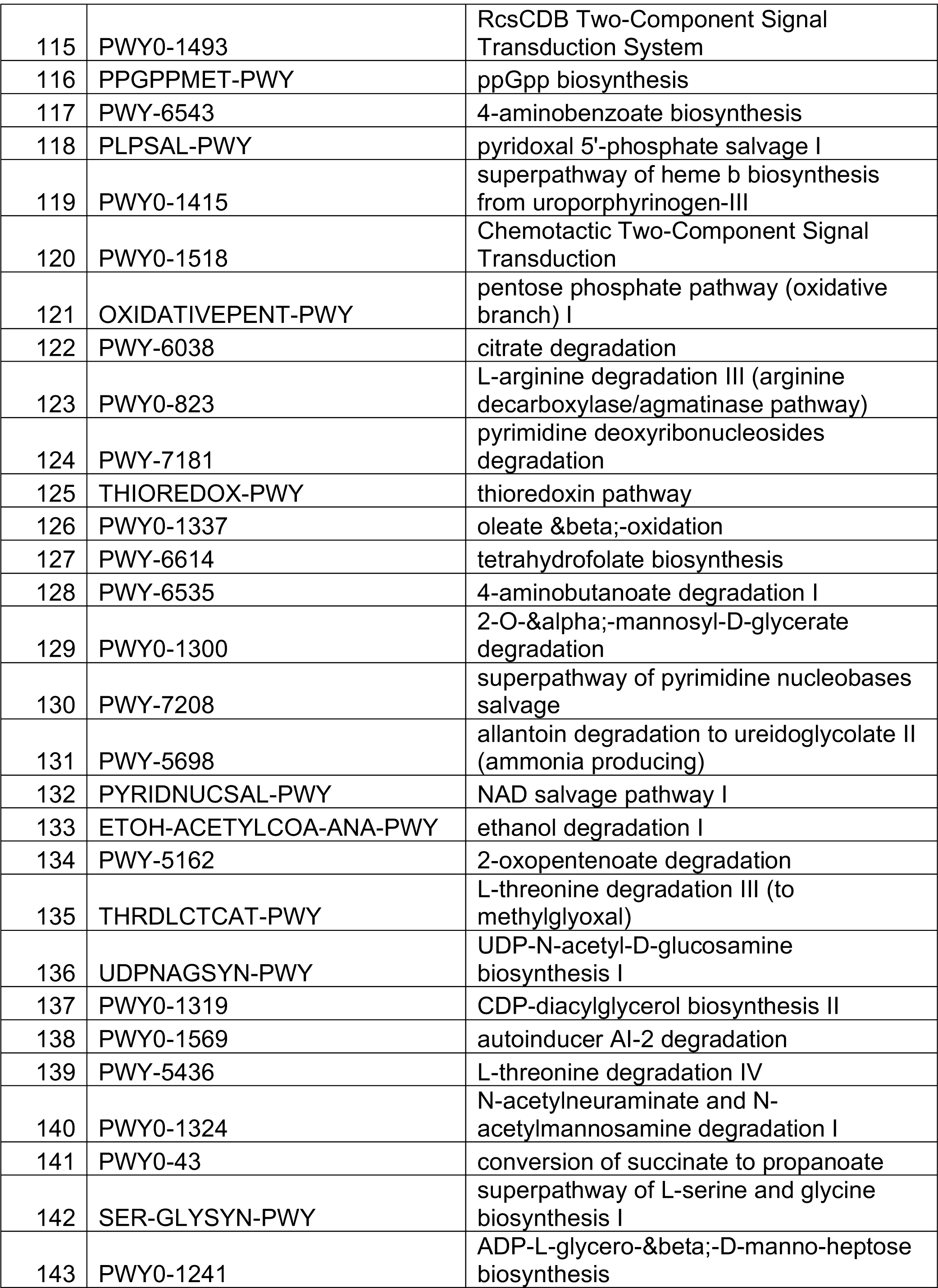
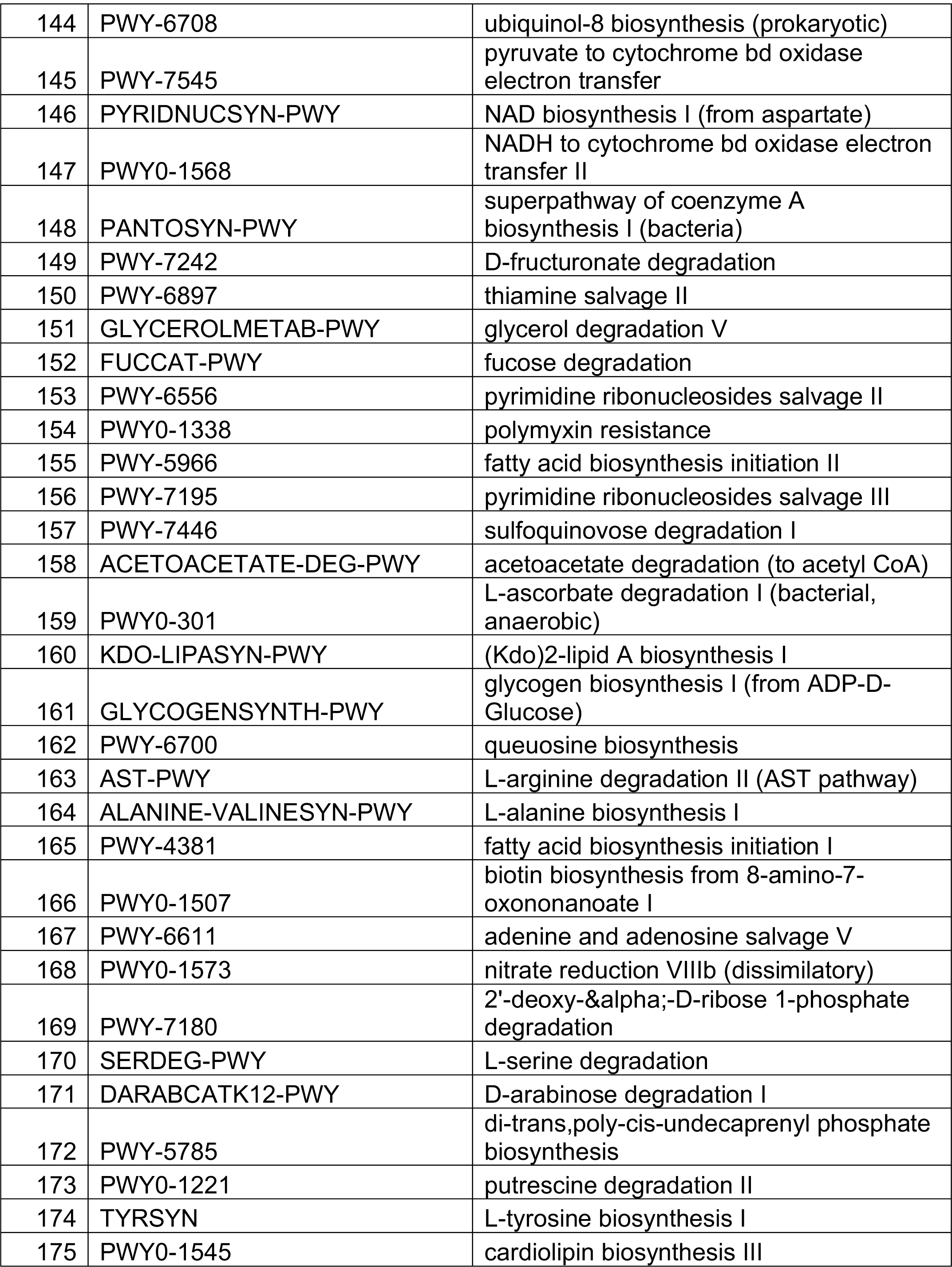
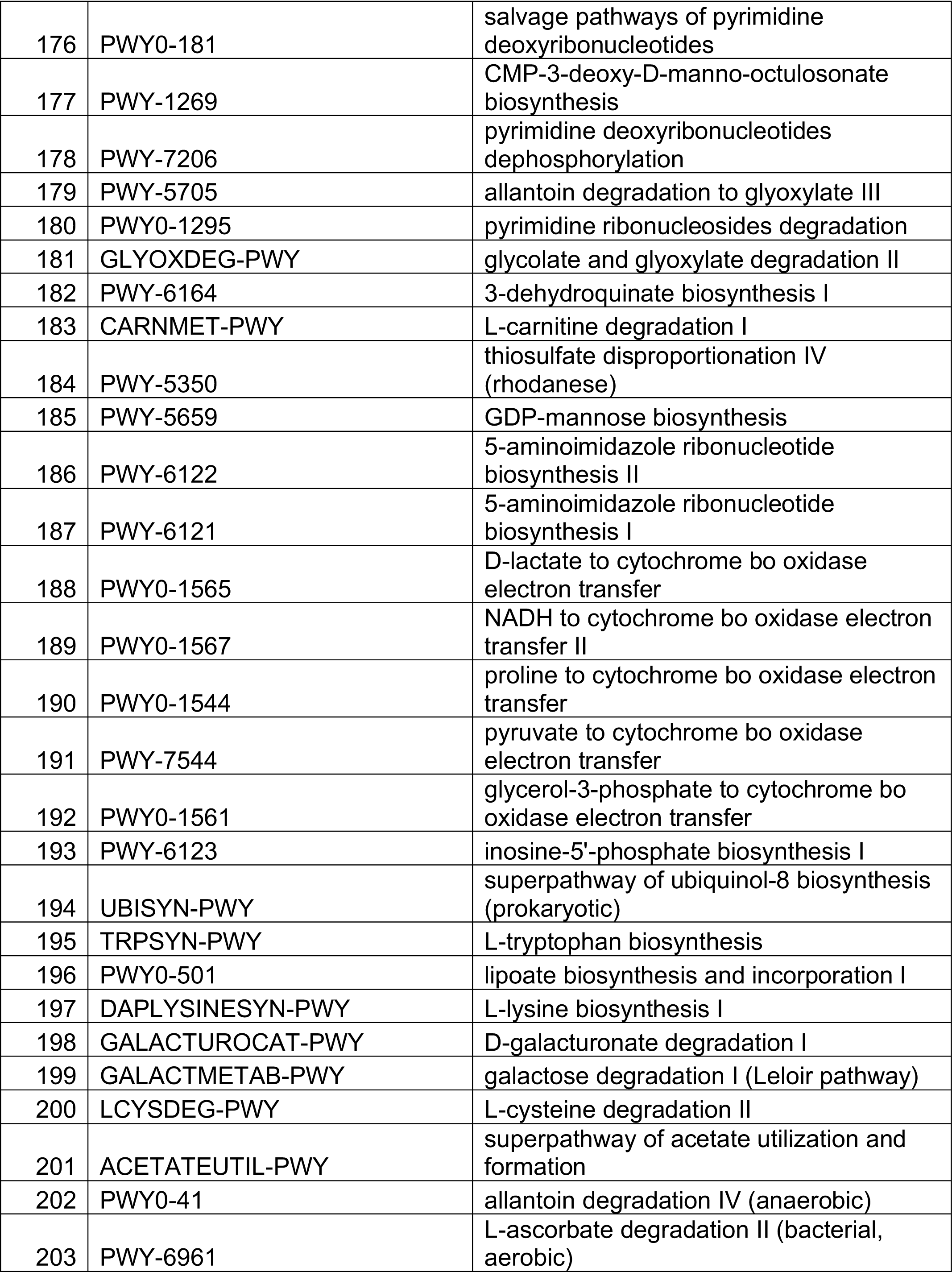
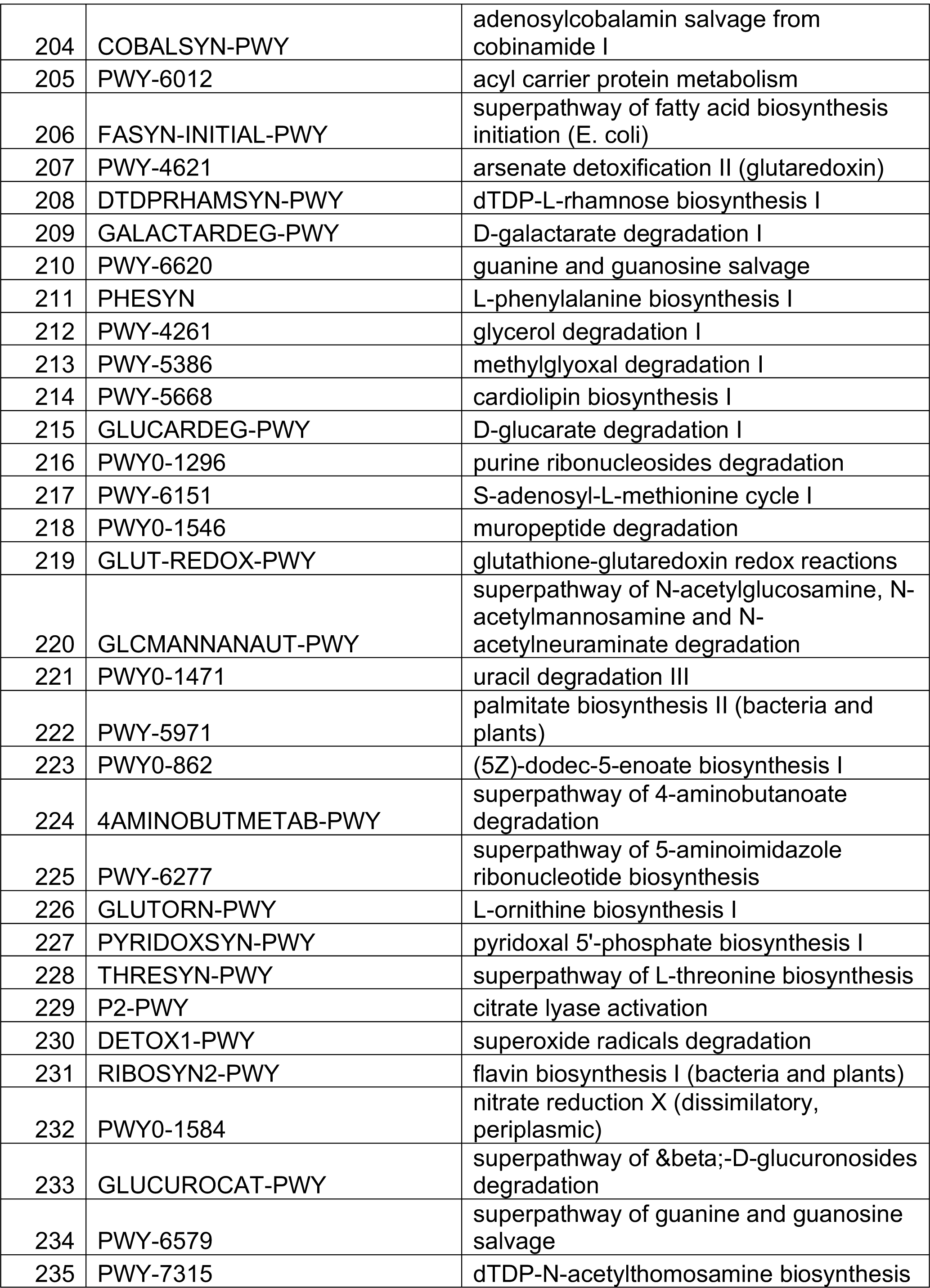
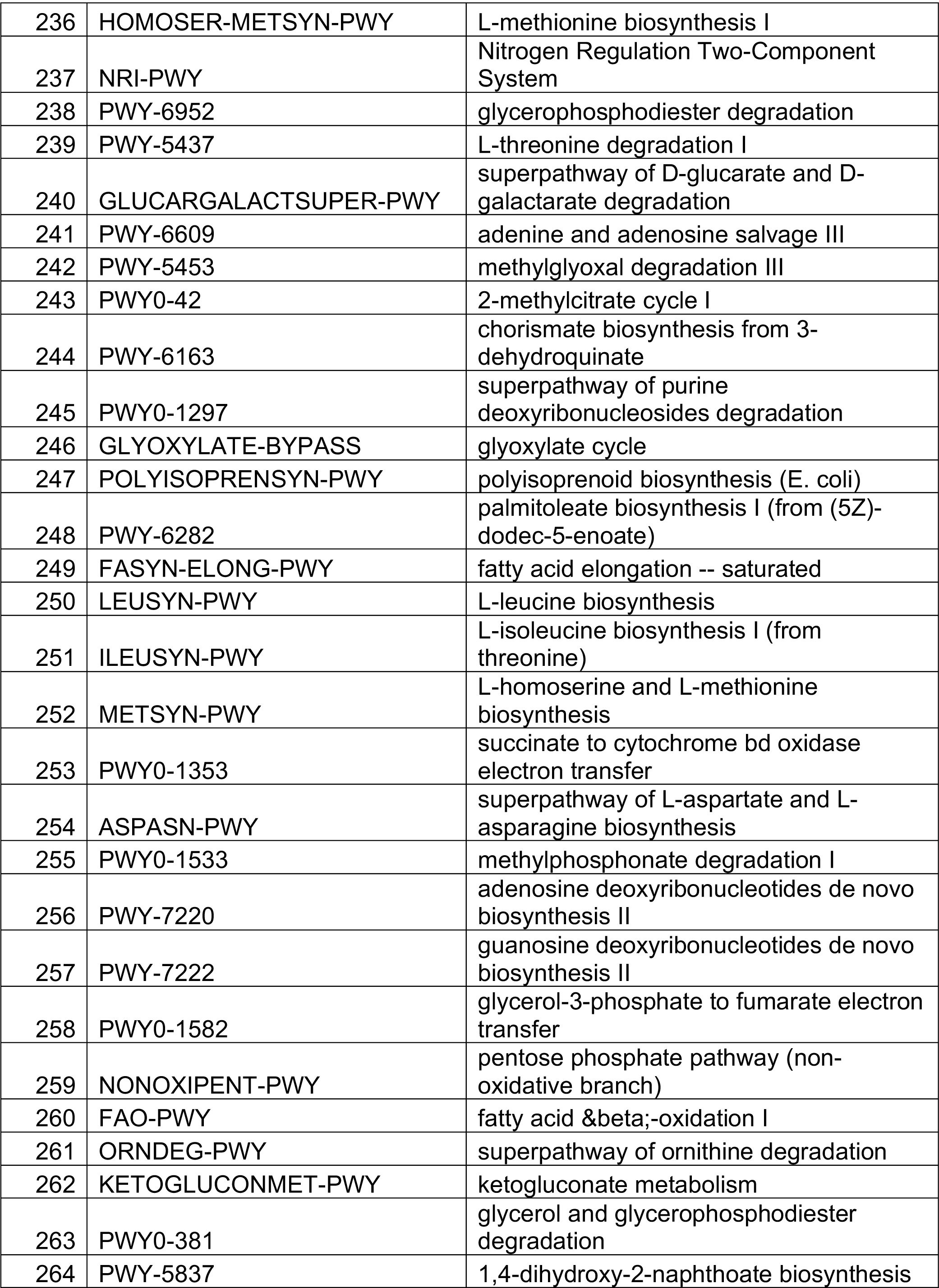
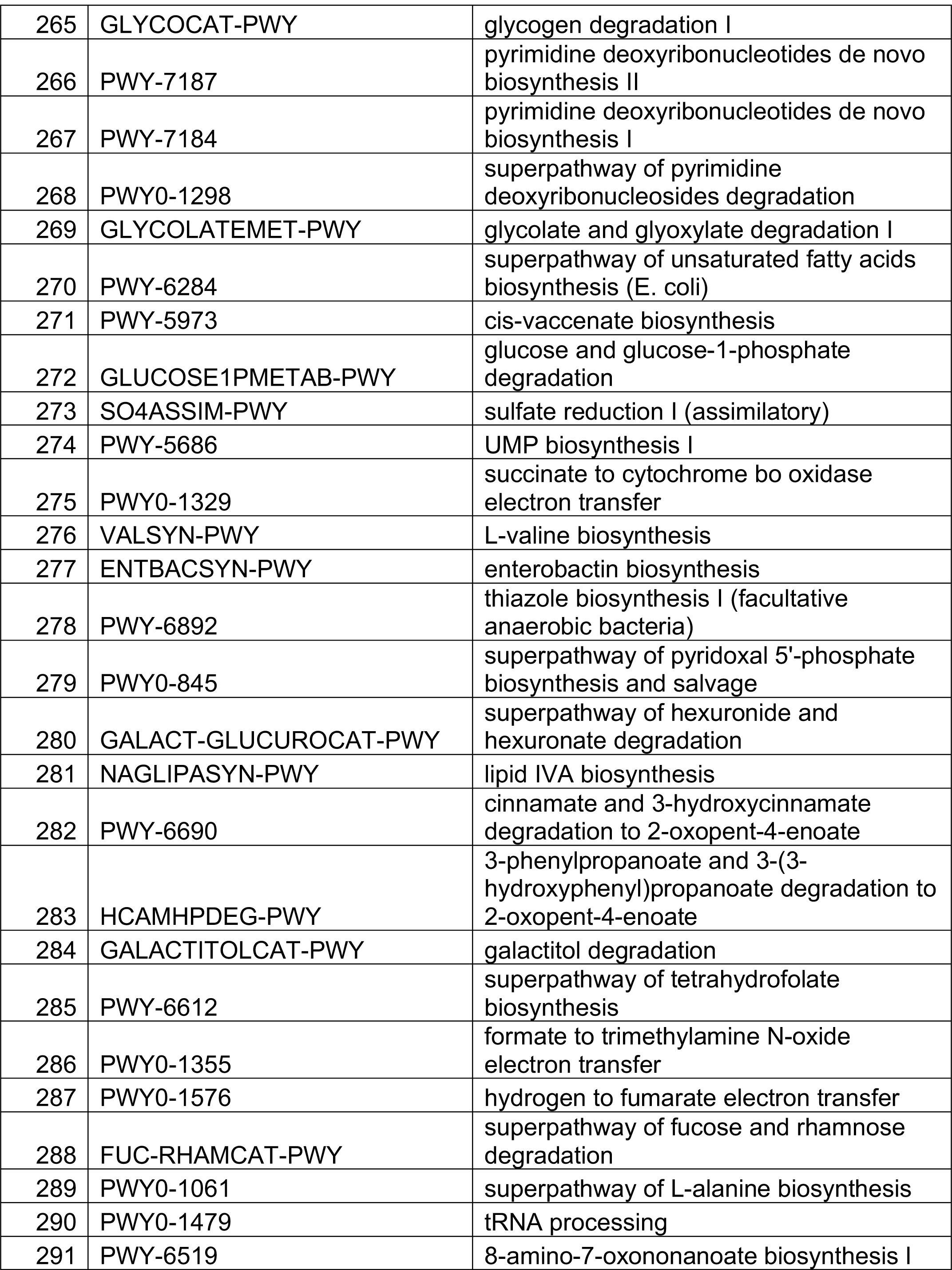
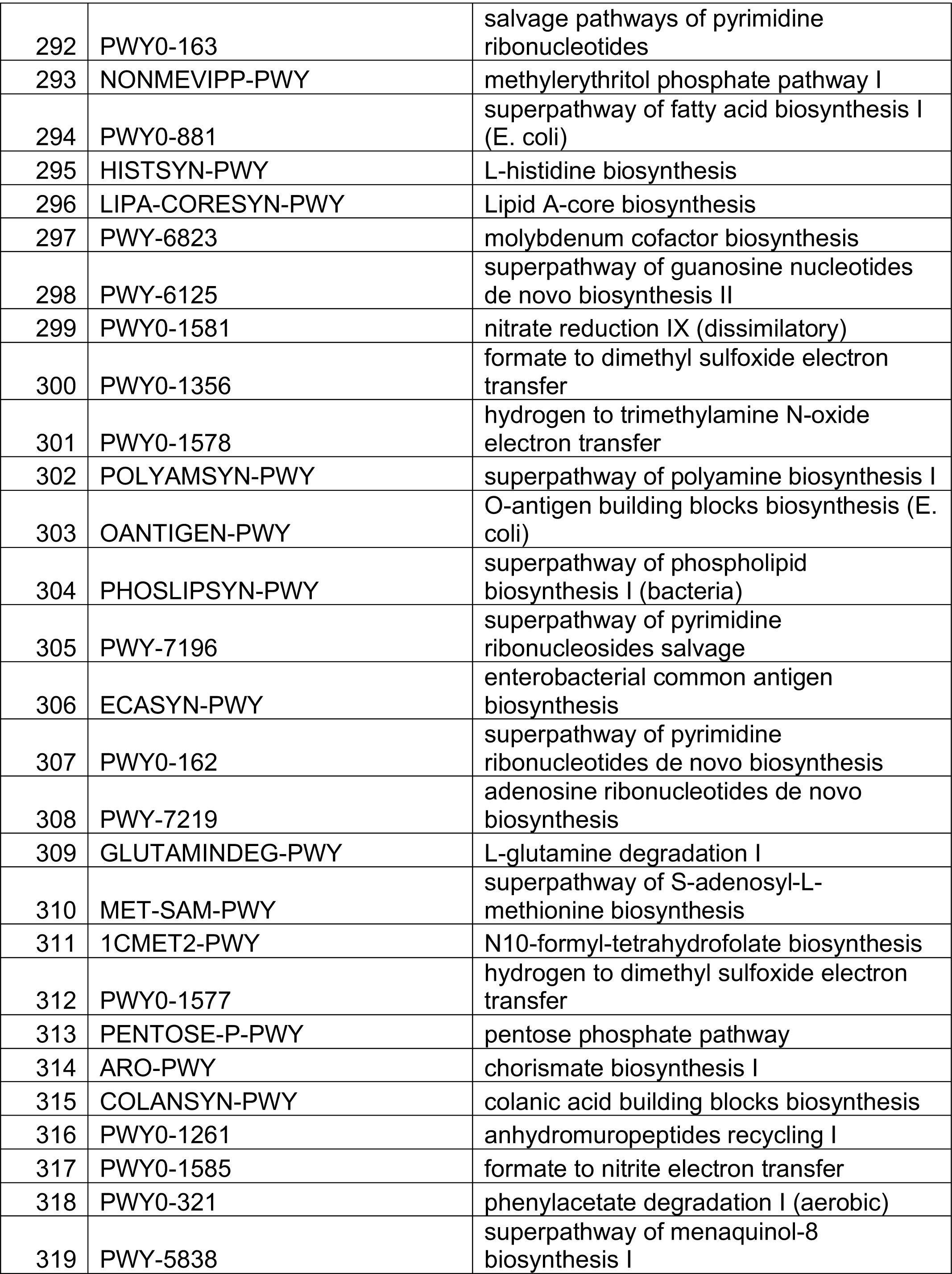
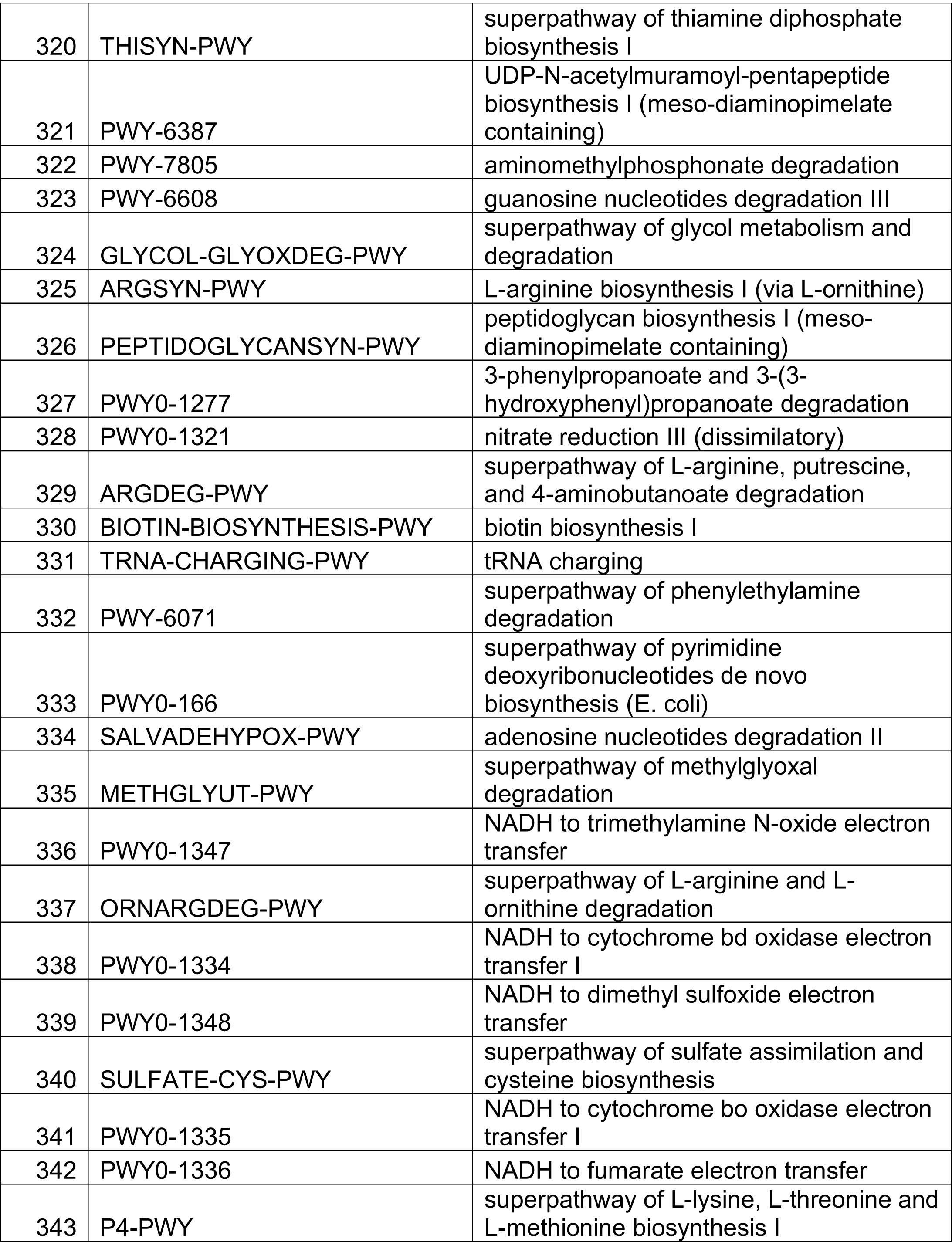
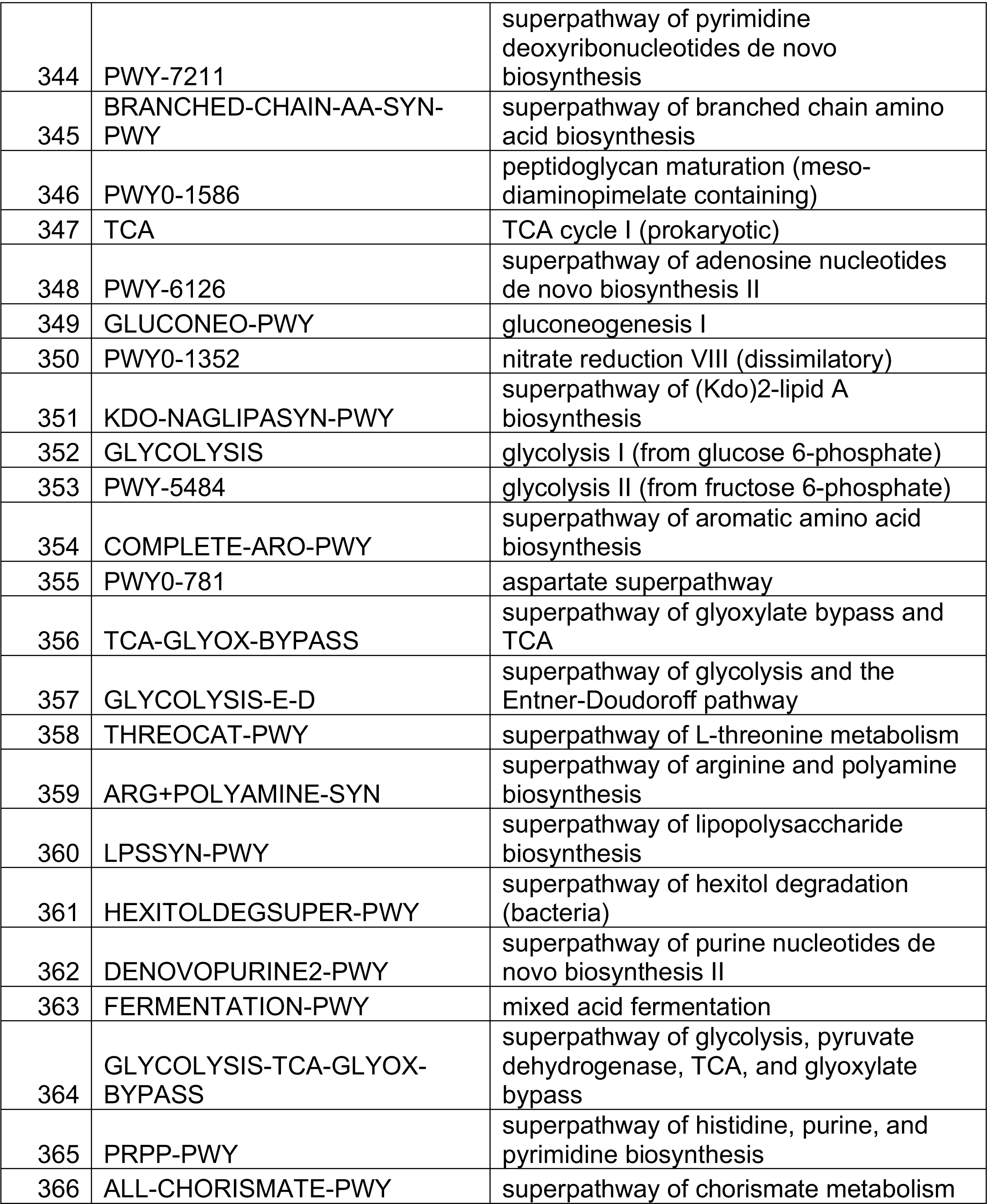
EcoCyc Pathways IDs and name of pathway for the labels used in Supplemental Figure S1.

**Supplemental Table S2.**
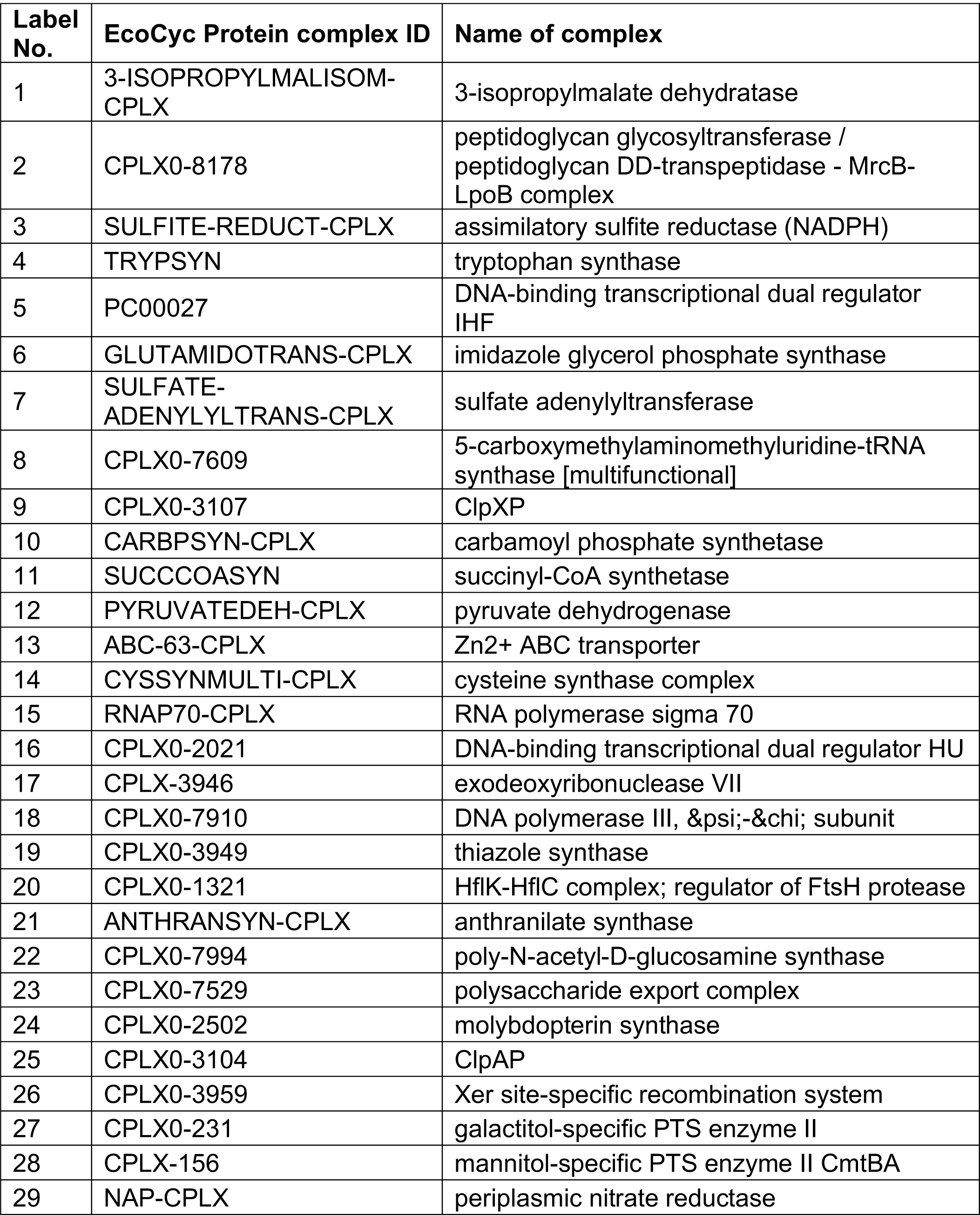
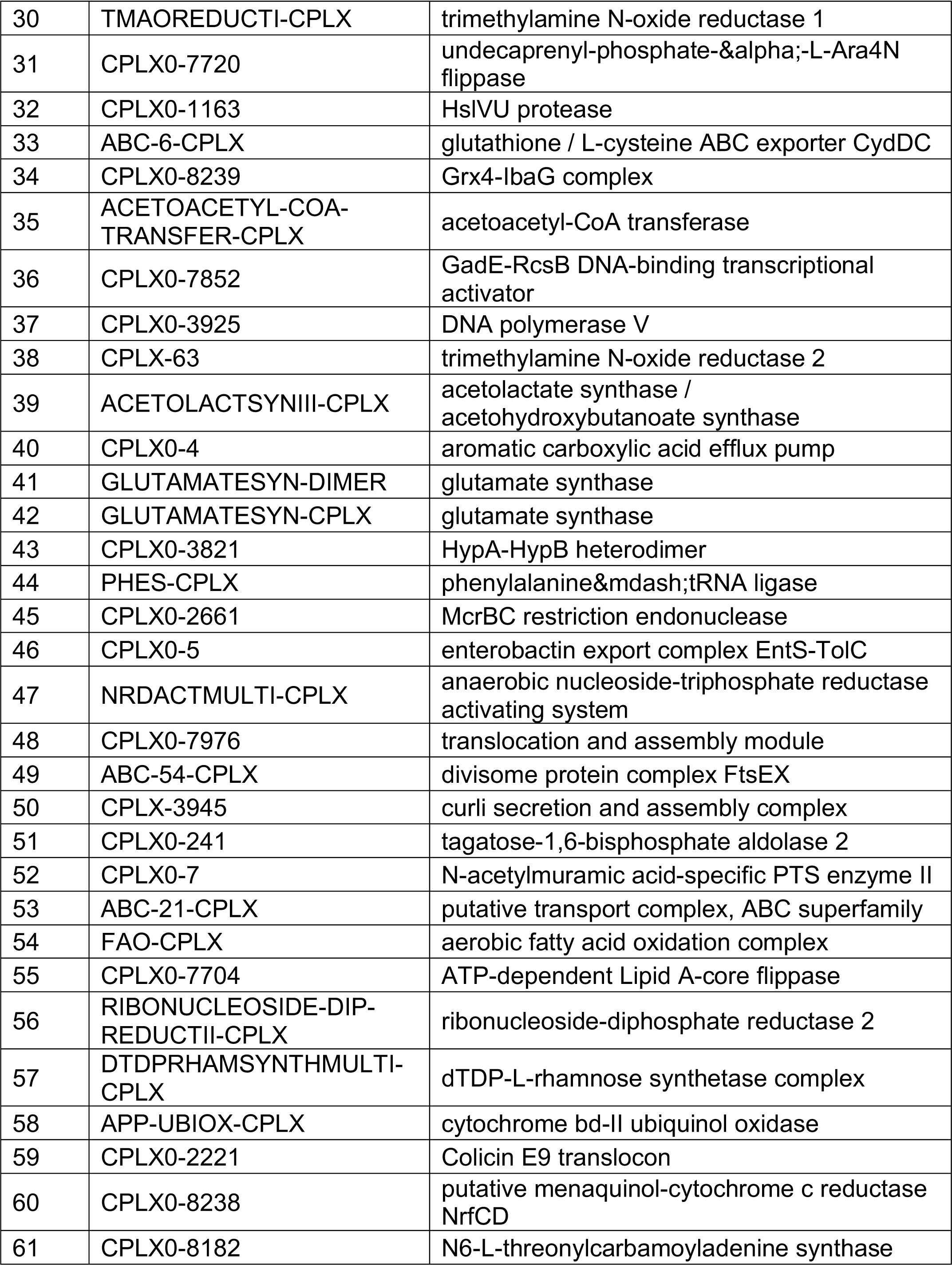
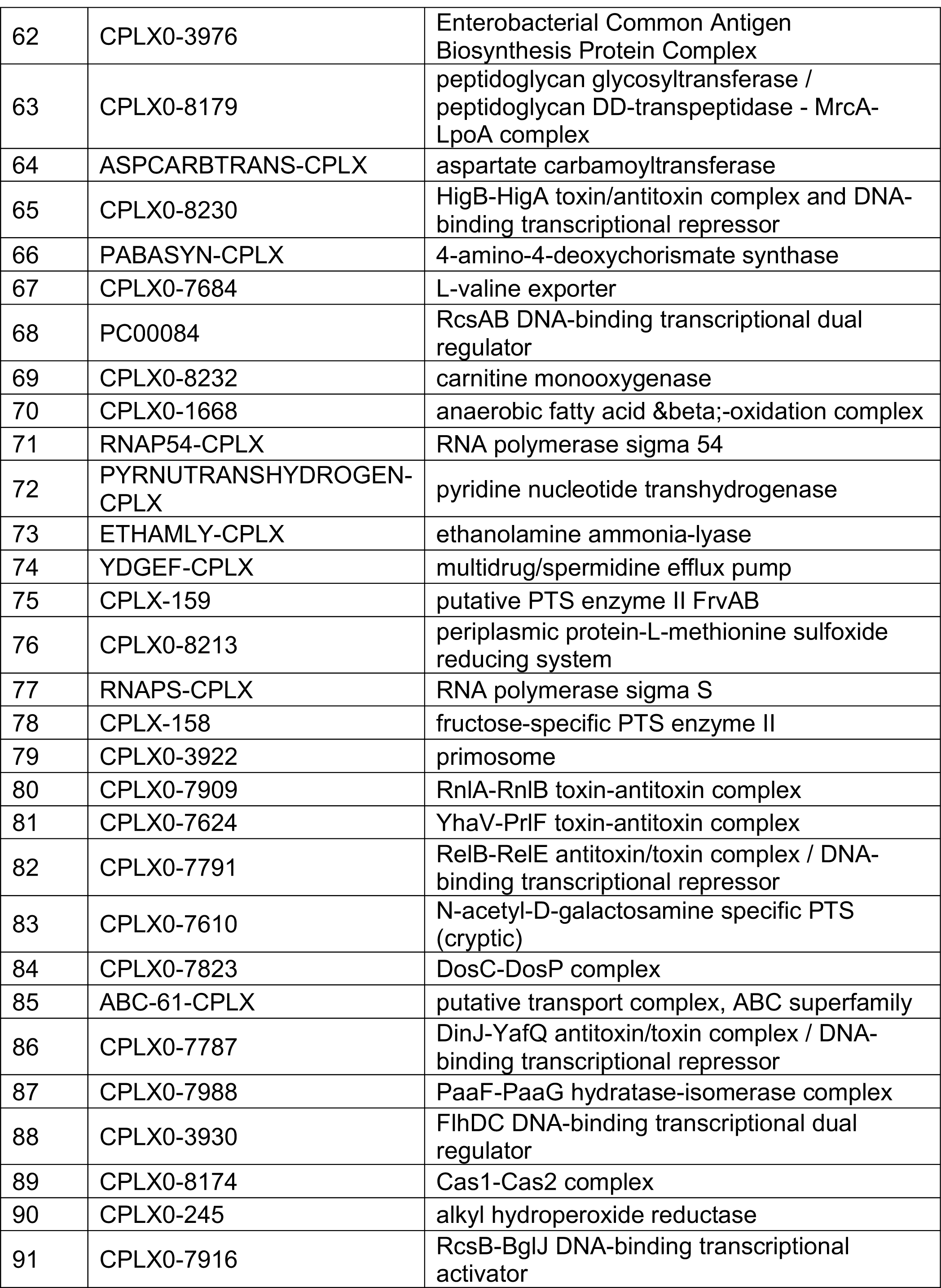
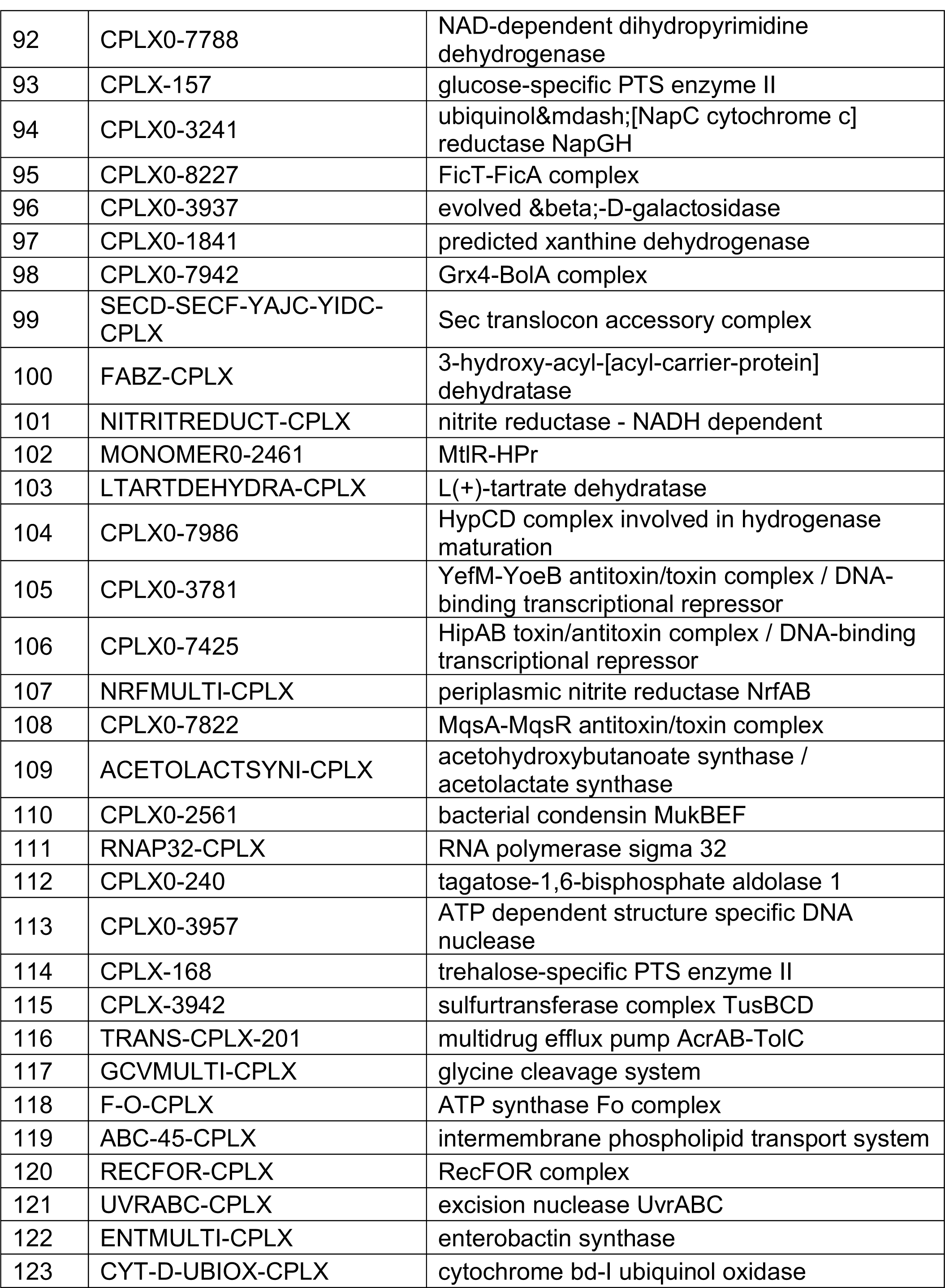
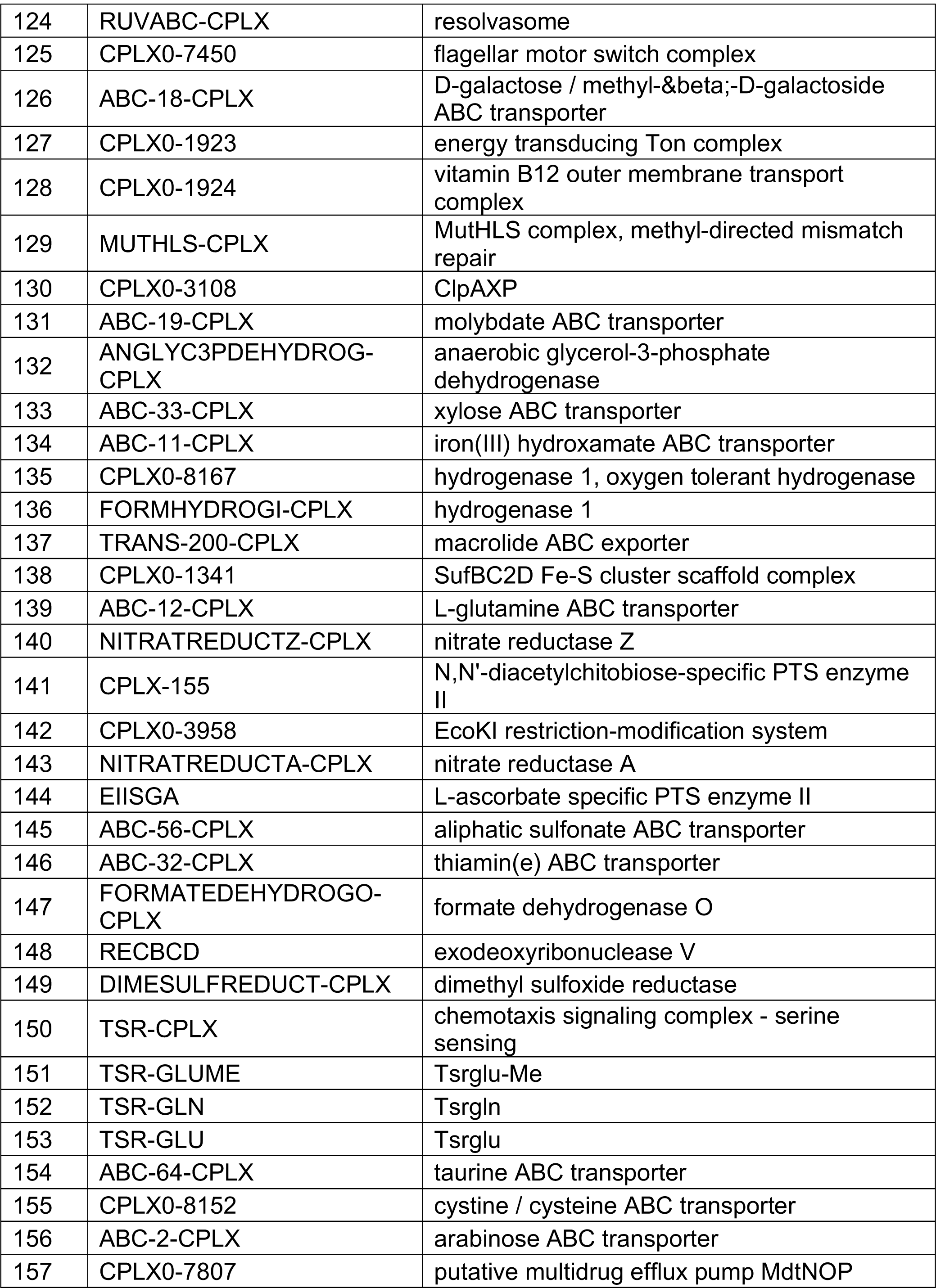
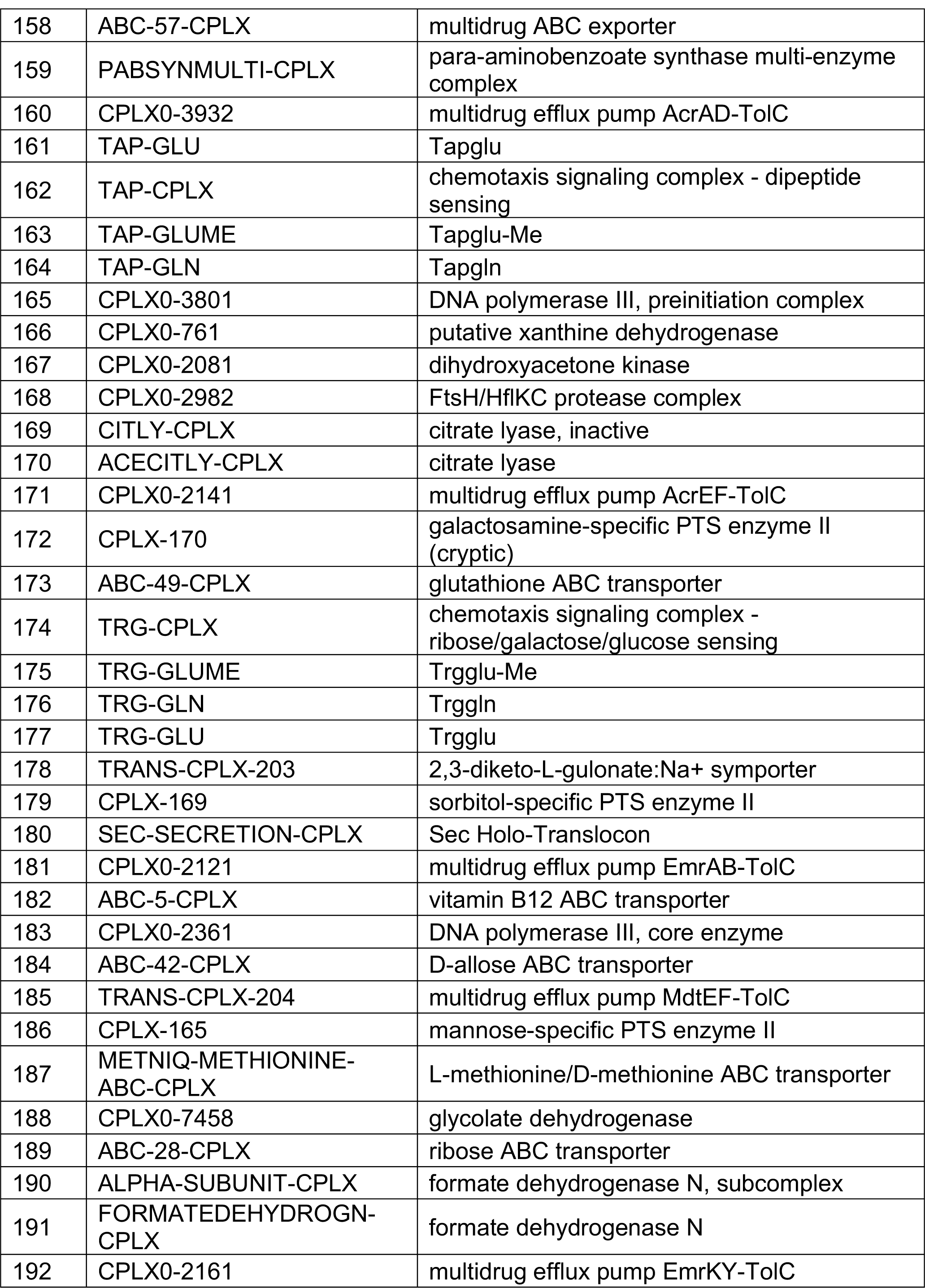
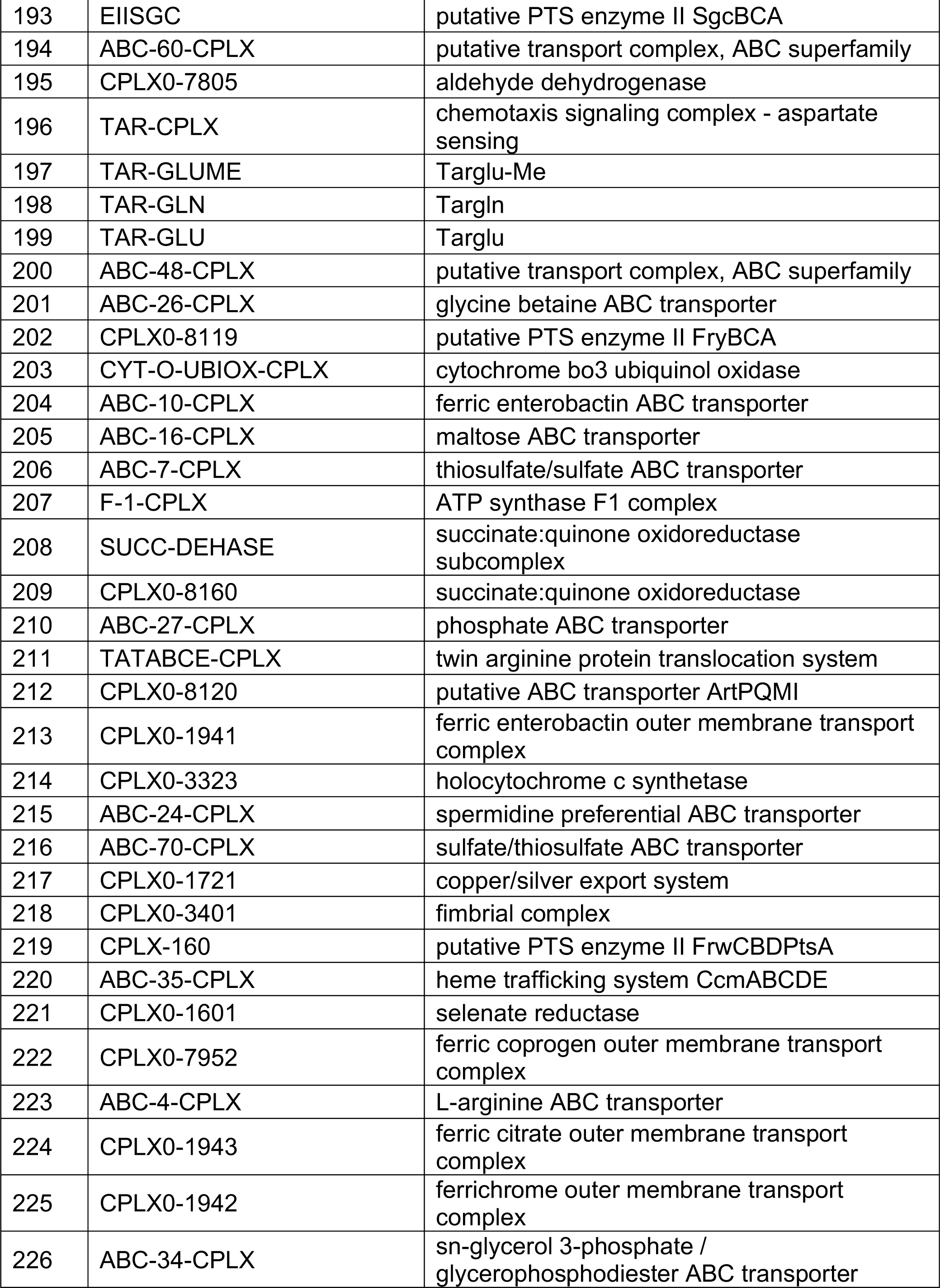
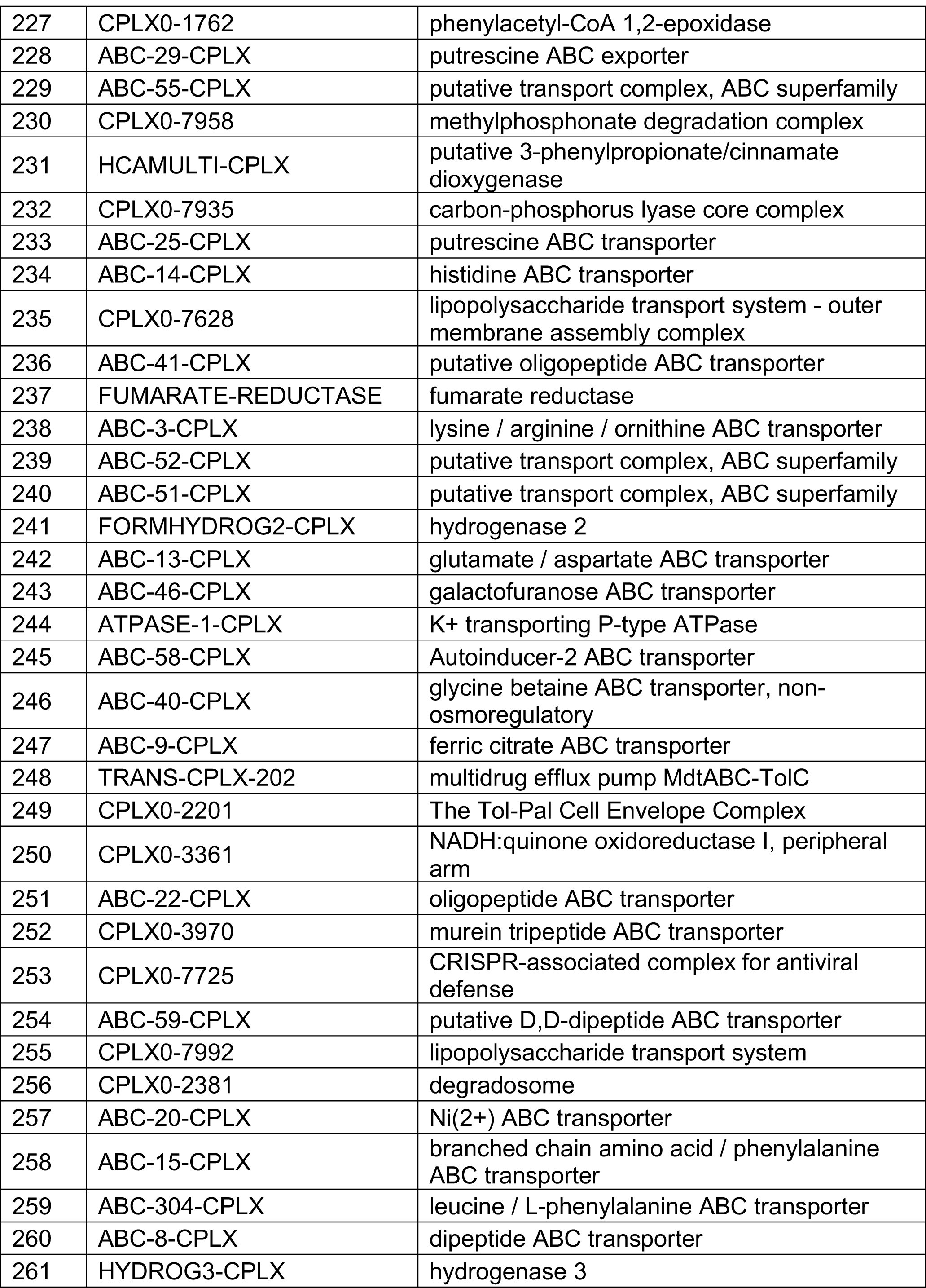
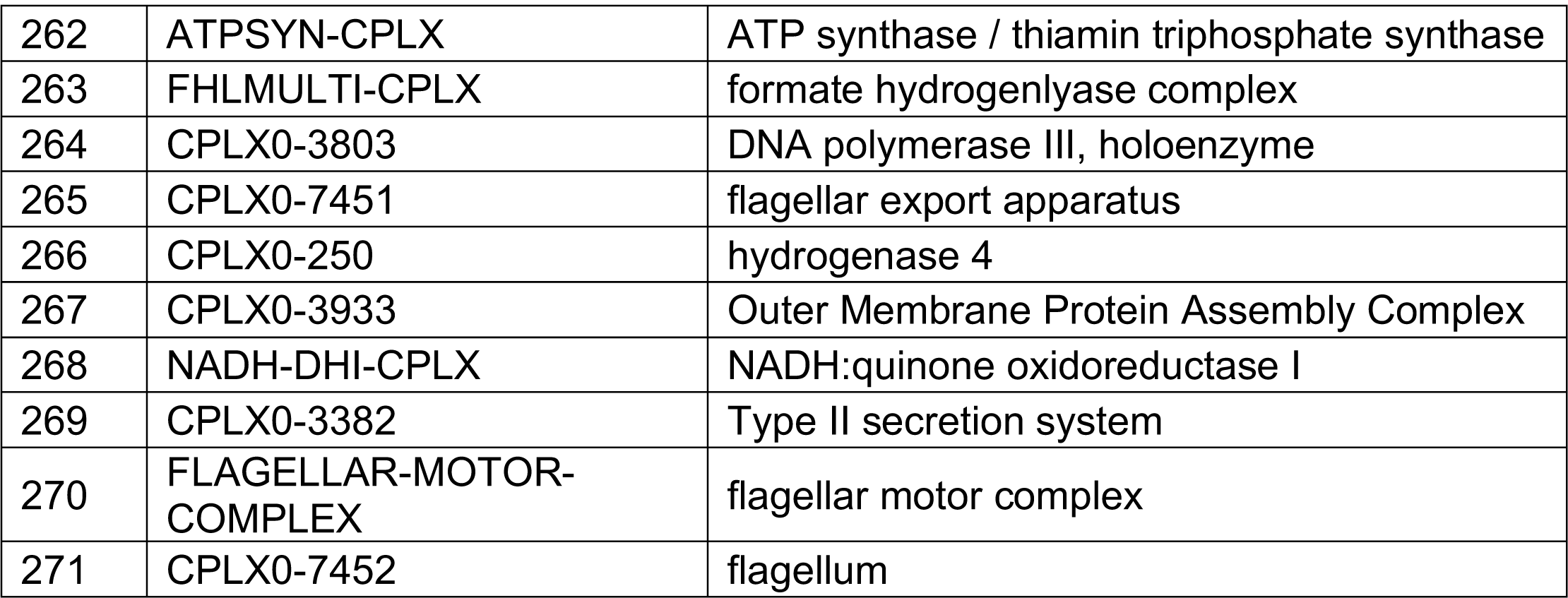
EcoCyc protein complex IDs and name of protein complex for the labels used in Supplemental Figure S2.

**Supplemental Figure S1.**
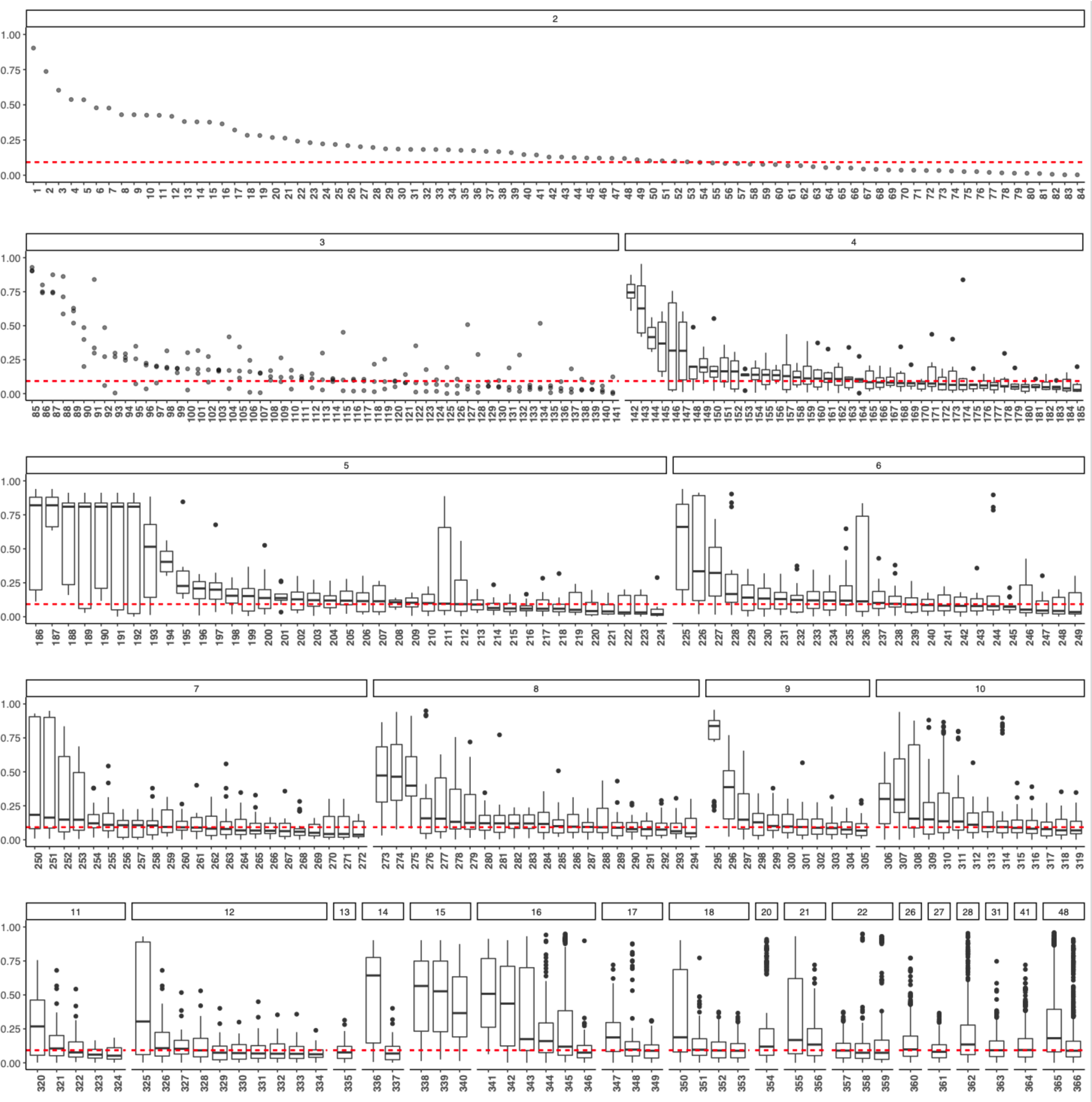
Phenotypic profile similarity for genes in the same heteromeric protein complex. The distribution of phenotypic profile similarity values determined by |PCC| for all pairwise combinations of genes assigned to the same EcoCyc pathway. In the figure, the pathways are sorted by (i) the number of genes in the pathway and then (ii) the median |PCC| value. The names of the pathways are indicated by numeric labels, which are defined in supplemental Table S1. The dashed line shows the average |PCC| value for random pairs of genes. For pathways that have two or three members, the results are shown as scatter plots. For pathways with more than three genes, the results are shown as box plots with the outliers shown as black dots.

**Supplemental Figure S2.**
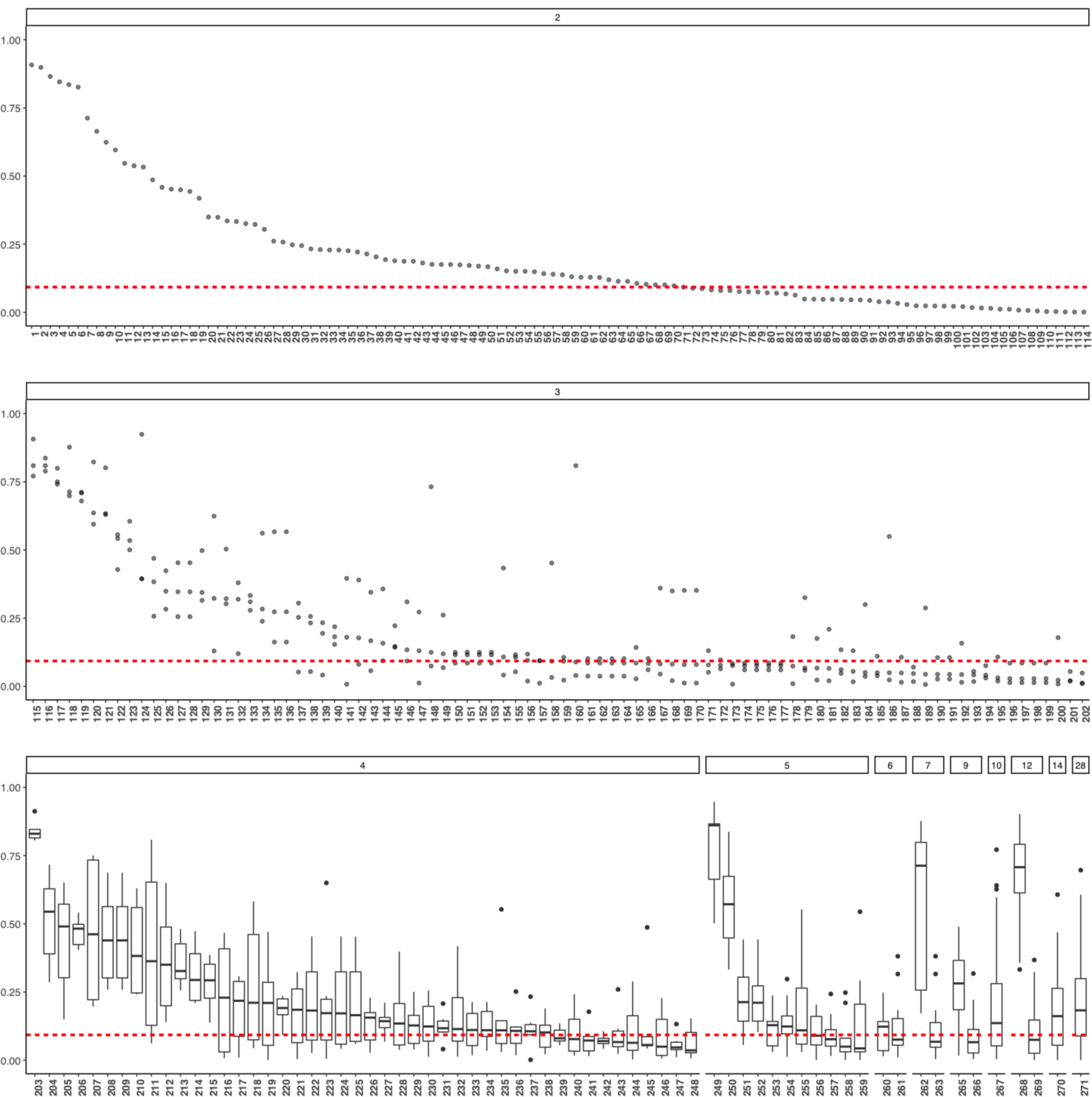
Phenotypic profile similarity for genes in the same EcoCyc heteromeric protein complex. The distribution of phenotypic profile similarity values determined by |PCC| for all pairwise combinations of genes assigned to the same EcoCyc heteromeric protein complex. In the figure, the pathways are sorted by (i) the number of genes in the complex and then (ii) the median |PCC| value. The names of the complexes are indicated by numeric labels, which are defined in supplemental Table S2. The dashed line shows the average |PCC| value for random pairs of genes. For protein complexes that have two or three members, the results are shown as scatter plots. For protein complexes with more than three genes, the results are shown as box plots with the outliers shown as black dots.

**Supplemental Figure S3.**
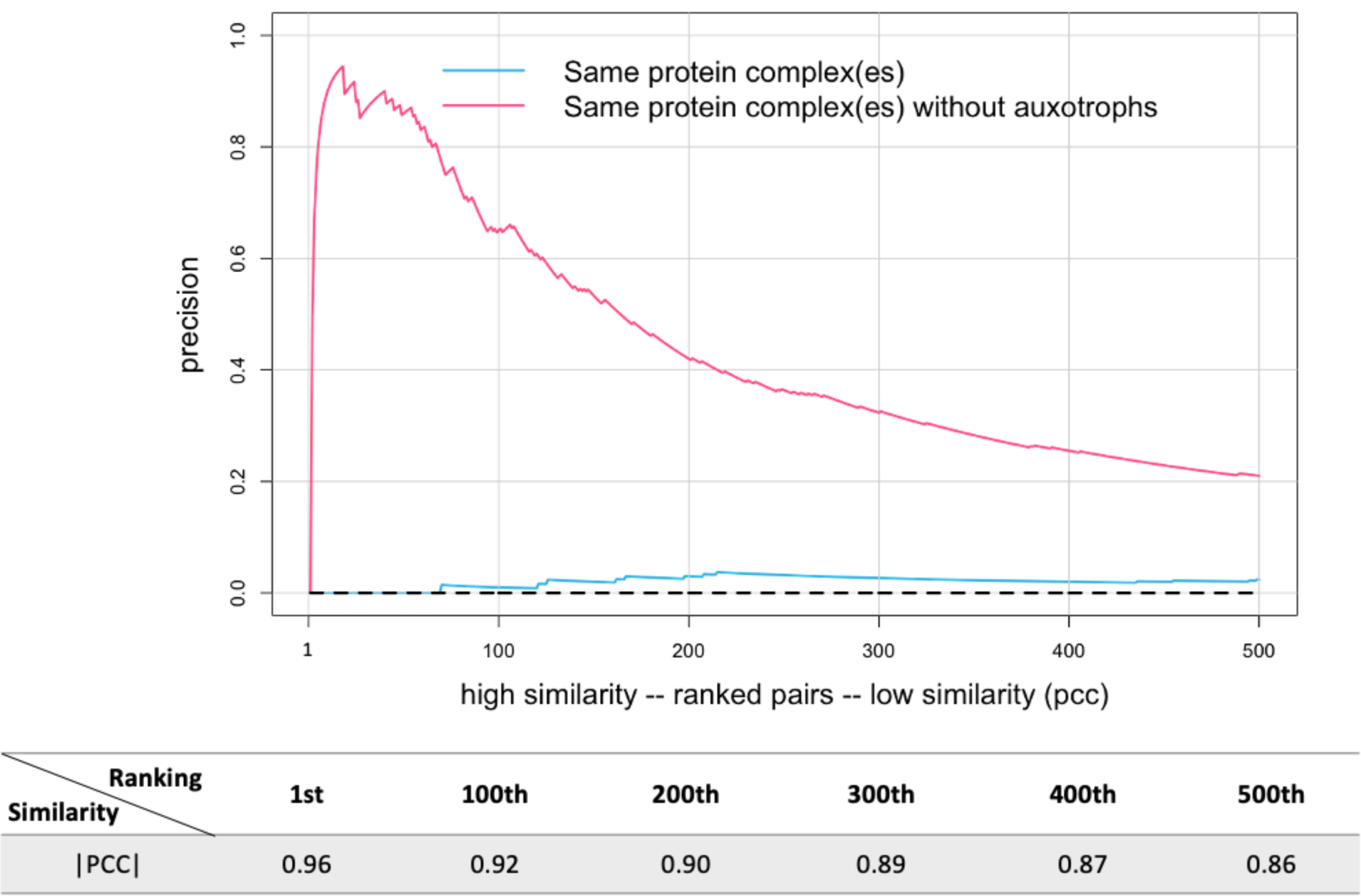
Precision increased when auxotrophic mutants were excluded. Gene pairs were ranked from high to low similarity based on |PCC| and plotted versus precision, calculated as described in the text (only the first 500 gene pairs are shown). The dashed line shows precision for randomly ordered gene pairs (negative control). The correspondence between phenotypic profile similarity based on |PCC| and ranking is shown below the graph.

**Supplemental Figure S4.**
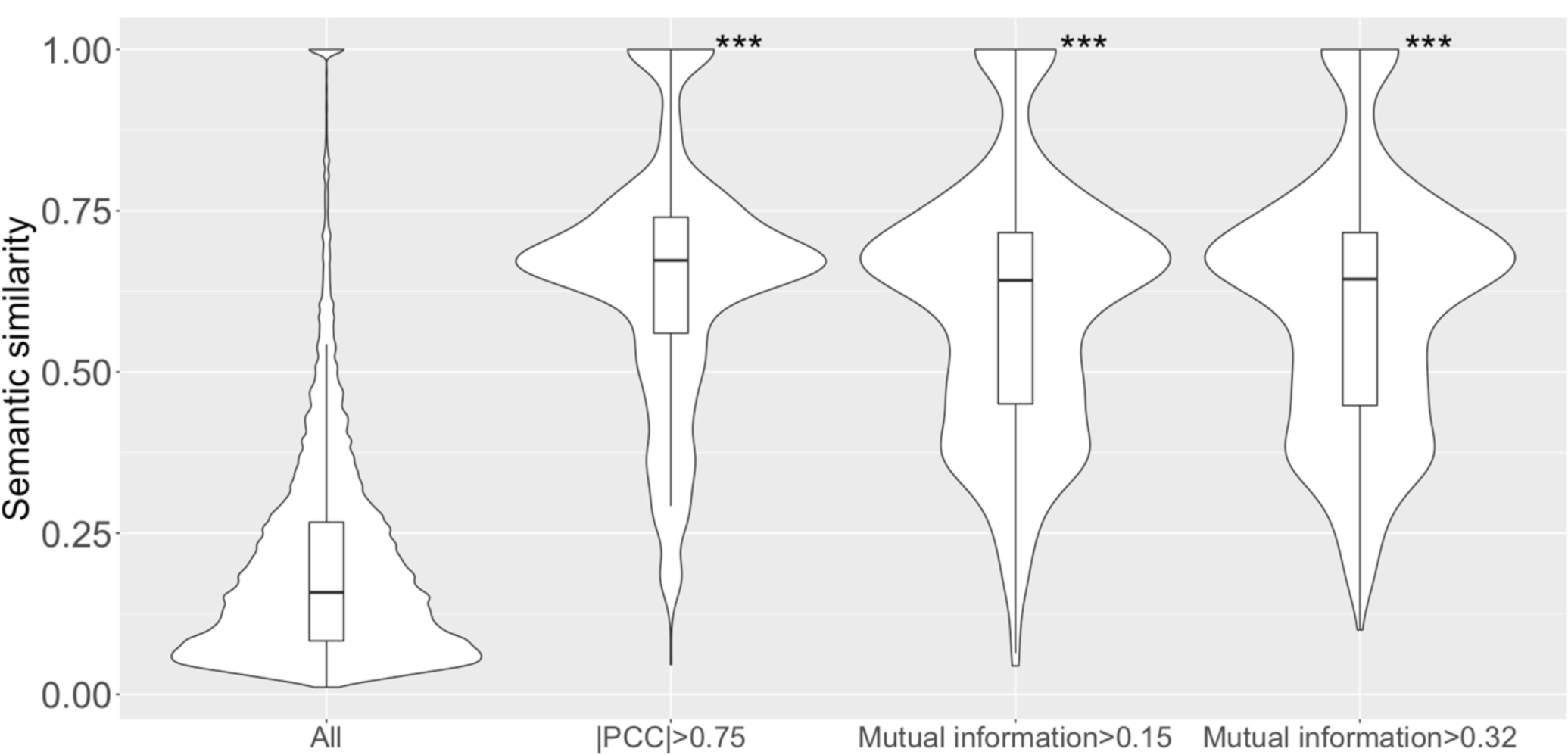
Higher semantic similarity and phenotypic profile similarity were still found when GO biological process annotations with an IEA evidence code were excluded. Violin plots of semantic similarity for, from left to right: all gene pairs annotated with GO biological process term(s); the subset of gene pairs with |PCC| >0.75; the subset of gene pairs with MI >0.15 (calculated based on qualitative fitness scores for all growth conditions); and MI >0.32 (calculated based on qualitative fitness scores for the collapsed set of growth conditions). The cutoffs of MI >0.15 for the third violin plot and MI >0.32 for the fourth violin plot were chosen so that all three subsets of gene pairs would contain the same number (∼1,000) of top-ranked gene pairs. ***: p-value <0.001 was determined by 1-sided Mann-Whitney U test, compared to all gene pairs.

